# Assembly of the essential SpoIVA coat protein at the surface of developing bacterial spores

**DOI:** 10.64898/2026.07.09.737436

**Authors:** Elda Bauda, Betty Fekade, Laure Bellard, Benoit Gallet, Seraphine Degroux, Emmanuelle Neumann, Caroline Mas, Kaitlyn Coleman, Aline Leroy, Gregory Effantin, Daphna Fenel, Jana Moravcova, Jiri Novacek, Christine Moriscot, Guy Schoehn, Christopher D.A. Rodrigues, Cecile Morlot

**Affiliations:** Univ. Grenoble Alpes, CNRS, CEA, IBS, F-38000 Grenoble, France; School of Life Sciences, University of Warwick, Coventry, UK; Integrated Structural Biology Grenoble (ISBG), UAR 3518, CNRS, CEA, Univ. Grenoble Alpes, EMBL, F-38042, Grenoble, France; CEITEC-Central European Institute of Technology, Masaryk University, 62500 Brno, Czech Republic

**Keywords:** Cryo-FIB/SEM, cryo-electron tomography, cellular electron microscopy, sporulation, coat proteins, SpoIVA, *Bacillus subtilis*

## Abstract

Bacterial spores owe their remarkable resistance properties to a multilayered coat, one of the most resilient and durable biological structures on Earth. Assembled at the surface of the outer forespore membrane, the coat comprises dozens of proteins organized into distinct layers. Its formation is initiated by SpoIVA, which is proposed to form a polymeric scaffold for the innermost coat layer. Although SpoIVA has been shown to polymerize into filaments *in vitro*, there is currently no evidence demonstrating the formation of such assemblies *in vivo*, and the mechanism underlying its oligomerization remains unresolved. In this study, cryo-focused ion beam milling combined with cryo-electron tomography of sporulating *Bacillus subtilis* cells reveals that the SpoIVA layer consists of polymers that form track-like structures radiating from the mother cell-proximal forespore pole and extending directionally toward the distal pole. Subtomogram averaging further sheds light on their organized architecture, harboring a straight orientation, uniform spacing, and embedding in the outer forespore membrane. These observations also define SpoIVA spatial orientation relative to the outer forespore membrane. Furthermore, AlphaFold3 predictions, combined with biophysical and functional assays, show that SpoIVA dimerizes through its central and C-terminal regions. We further show that dimerization promotes SpoIVA localization around the forespore but is dispensable for polymer formation, which relies on the ATPase domain. Altogether, these findings suggest a dual oligomerization mechanism, in which SpoIVA transitions from dimers to linear track-like polymers, and reveal that these assemblies play critical roles in coat assembly and spore development.

## Introduction

Bacterial spores are differentiated cells whose unique cellular structures enable them to survive under extreme conditions, including high temperatures, detergents and irradiation (Setlow and Christie, 2023). Metabolically dormant, spores are also impervious to antibiotics. Such properties are valuable in the context of spore-based therapies, but cause a major burden of dissemination and persistence in the case of spore-forming pathogens (McKenney et al., 2013; Setlow and Christie, 2023). Understanding how the cellular structures responsible for spore resistance assemble at the nano- and ultrastructural level is therefore crucial for developing strategies to either harness or counteract spore properties. In this study, we present the first cellular structural analysis of the large oligomers formed by SpoIVA, the sole sporulation protein conserved in all spore-formers (Galperin et al., 2022), whose supramolecular assembly underpins the remarkable resilience of bacterial spores.

Sporulation is a complex developmental process found in Bacillales and Clostridia within the Firmicutes phylum. It relies on a sophisticated differentiation process leading to the release of a mature spore into the environment. At the onset of sporulation, asymmetric division generates a sporangium composed of a large mother cell and a smaller forespore (Tan and Ramamurthi, 2014). These two cells then undergo dramatic morphological changes, orchestrated by compartmentalized gene expression programs (Hilbert and Piggot, 2004). Following asymmetric division, the mother cell membrane expands around the forespore during a phagocytic-like phase called “engulfment”. The engulfed forespore becomes surrounded by its cytoplasmic membrane, called the inner forespore membrane (IFM), and the outer forespore membrane (OFM), inherited from the mother cell. These two membranes define a space called the intermembrane space (IMS). Engulfment is coordinated with the assembly of two protective envelope structures called the cortex and the coat (Driks and Eichenberger, 2016; Popham and Bernhards, 2015). The cortex, a modified peptidoglycan synthesized in the IMS, maintains the spore in a dehydrated state, protecting its cytoplasmic content and preserving dormancy (Popham and Bernhards, 2015). The coat, a highly resistant integument composed of over 80 different proteins, assembles into multiple layers with distinct dimensions, organization, and properties (Bauda et al., 2024; Driks and Eichenberger, 2016; McKenney et al., 2013; McKenney and Eichenberger, 2012). In *Bacillus subtilis*, these layers include the basement layer, the inner coat, the outer coat and the crust.

Coat assembly is controlled by morphogenetic proteins, which establish a hub of interactions with coat proteins to guide the deposition of the distinct layers. This process is initiated by the interdependent localization of SpoVM and SpoIVA at the mother cell-forespore interface (Ramamurthi et al., 2006). These proteins, which localize around the forespore, are essential for coat morphogenesis and spore resistance; in their absence, coat layers detach from the OFM, forming swirls in the mother cell cytoplasm (Bauda et al., 2024; Catalano et al., 2001; Ramamurthi et al., 2006). SpoIVA is the only universally conserved morphogenetic protein (Galperin et al., 2022). *In vitro*, it forms filaments whose polymerization and stability are dependent on ATP binding and hydrolysis (Castaing et al., 2013; Peluso et al., 2019; Ramamurthi and Losick, 2008). *In vivo*, the layer formed by SpoIVA has been visualized as an organized structure at the surface of the OFM, but its architecture has yet to be elucidated (Bauda et al., 2024). SpoVM is a 26 amino-acids amphipatic protein with affinity toward positively curved membranes (Ramamurthi and Losick, 2009). The hydrophobic face of its α-helix is buried in the lipid bilayer, while its hydrophilic face interacts with the extreme C-terminus of SpoIVA (Gill et al., 2015). SpoIVA also directly interacts with two other morphogenetic proteins, SpoVID (via its C-terminal region) and SafA (through an undefined region), both of which are required for the formation of the basement and inner coat layers (Bauda et al., 2024; Delerue et al., 2022; Müllerová et al., 2009; Wang et al., 2009). SpoVID harbors a N-terminal β-sandwich domain, and a C-terminal region containing a disordered loop and a LysM domain. SpoVID interacts preferentially with polymerized SpoIVA compared to unpolymerized SpoIVA, and its LysM domain binds the Lipid II peptidoglycan precursor (Delerue et al., 2022; Müllerová et al., 2009; Wang et al., 2009). These observations led to the hypothesis that SpoVID sequesters Lipid II to prevent cortex assembly until it binds SpoIVA polymers, ensuring orchestrated assembly of the spore envelope.

Despite its central role as the scaffold for coat layers, the molecular mechanism of SpoIVA oligomerization and the organization of the SpoIVA layer at the surface of the OFM remain unknown. In this work, using cryo-electron tomography (cryo-ET) on cell sections prepared by cryo-focused ion beam milling (cryo-FIB), AlphaFold3 predictions, biophysical analyses and functional assays, we uncover the architecture of SpoIVA track-like structures at the surface of *B. subtilis* forespores and provide insight into the mechanism of SpoIVA oligomerization.

## Results

### SpoIVA forms linear polymers at the surface of the forespore

To elucidate the architecture of the SpoIVA layer at the forespore surface, we performed cryo-FIB-ET on *B. subtilis* cells collected at 2 and 3 hours after their entry into sporulation. Tilt series were first acquired at medium magnification (pixel size of 4.5 Å) and high defocus to generate high-contrast tomograms, enabling visualization of SpoIVA distribution around the forespore. The quality of the cell lamellae allowed a particularly clear observation of the previously reported crenelated pattern, which emerged early during engulfment and progressively extended around the forespore as engulfment proceeded (Fig. 1A, T2) (Bauda et al., 2024). The most well-organized SpoIVA structures appeared after completion of engulfment, with an average height of 11.6 ± 0.5 nm (n = 70) and spacing of 2.4 ± 0.4 nm (n = 30) (Fig. 1A, T3).

**Figure 1.**
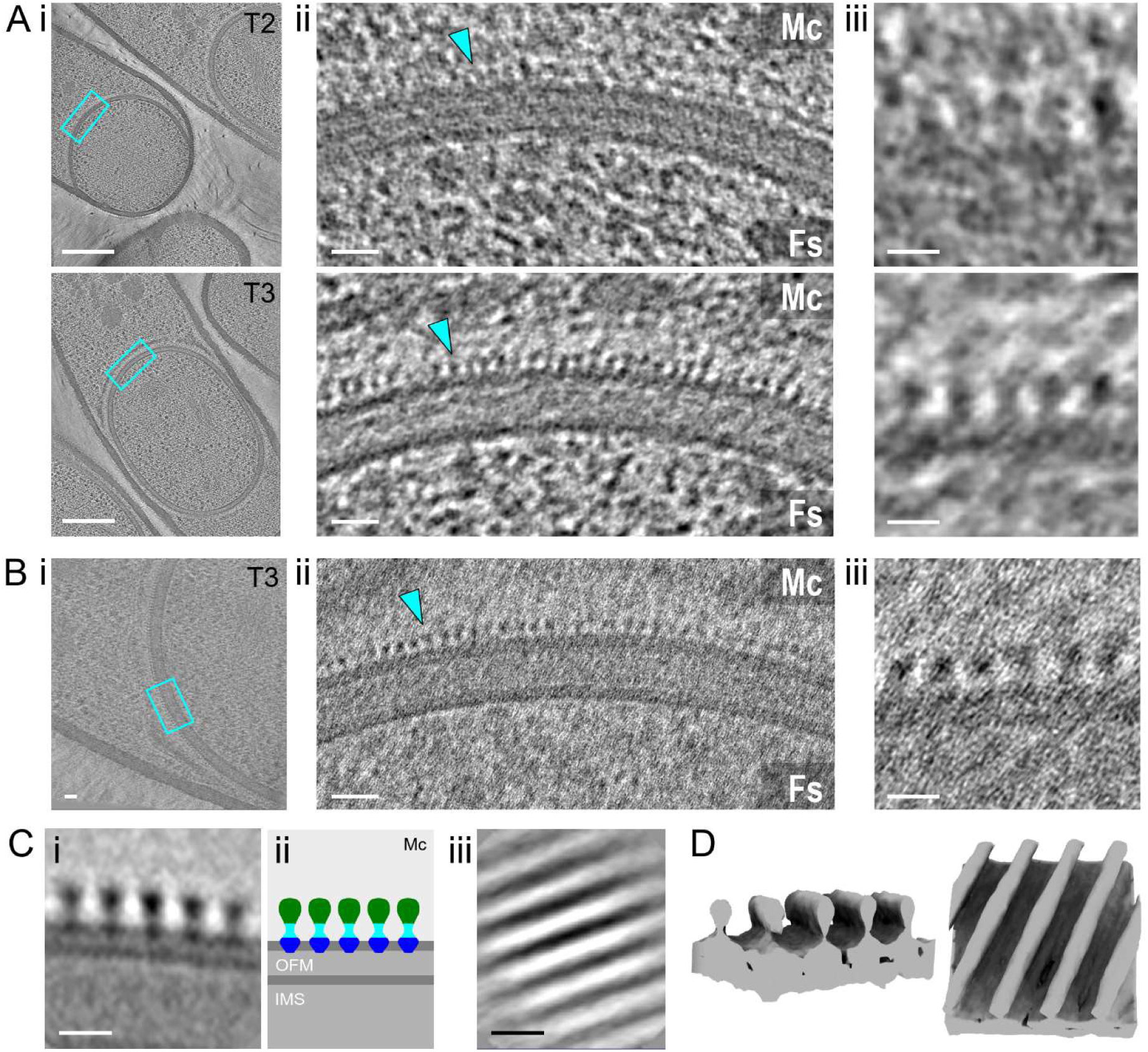
SpoIVA forms linear tracks at the forespore surface. **A-B.** Slices through cryo-electron tomograms of *B. subtilis* sporangia at different developmental stages, shown as full views (**i**; scale bars, 200 nm), and magnified views of the forespore-mother cell interface (**ii**; scale bars, 20 nm) and the outer forespore membrane (OFM) region (**iii**; scale bars, 10 nm). The crenelated SpoIVA layer (cyan arrowheads) is visible above the OFM. Cyan insets highlight the zoomed area. Tomograms were reconstructed from data acquired at medium magnification (pixel size 4.5 Å, high defocus) **(A)** or high magnification (pixel size 1.8 Å, low defocus) **(B)**. **C.** Subtomogram averaging of SpoIVA particles reveals linear, regularly spaced polymers at the forespore surface. Averaged structures are shown in two orientations (**i**, **iii**; scale bars, 10 nm), along with a schematic representation of the SpoIVA topology (**ii**). The tracks consist of a globular domain (green), a stalk region (cyan), and a base region (blue) embedded in the outer leaflet of the OFM. **D**. Surface representation of averaged SpoIVA tracks in oblique and top views. Fs, forespore; Mc, mother cell; IMS, intermembrane space.

To determine the *in situ* structure of the SpoIVA layer, we employed subtomogram averaging (STA), using tilt series collected at high magnification and low defocus (pixel size of 1.8 Å) from cryo-FIB lamellae. While these parameters provided high-resolution data, they resulted in a lower signal-to-noise ratio (Fig. 1B), complicating alignment and tomogram reconstruction. The small size of SpoIVA (∼ 55 kDa) further challenged particle picking. However, the high abundance of the protein around the forespore facilitated manual selection of 1,050 particles from 3 tomograms of T3 sporangia (Fig. 1B). Particle alignment and averaging revealed that SpoIVA forms linear structures aligned along the OFM (Fig. 1C-D). Within the reconstructed volume, these filament-like structures are parallel, uniformly spaced (2.2 ± 0.3 nm apart, n = 1,050), and exhibit a height of 9.10 ± 0.6 nm. Structurally, they feature a globular region (4.2 ± 0.3 nm in height, 5.0 ± 0.5 nm in width), positioned above a thinner and lower-density stalk (1.9 ± 0.3 nm in height), itself connected to a basal region (2.3 ± 0.2 nm in height, 4.4 ± 0.5 nm in width) embedded in the outer leaflet of the OFM (Fig. 1C-D). The continuity of the SpoIVA densities with the OFM suggests that they include SpoVM. Consistent with this and with SpoVM’s affinity for convex membranes (Ramamurthi et al., 2009; Wu et al., 2015), the crenelated pattern was also observed along positively curved regions of membrane vesicles in cells exhibiting membrane defects (Fig. S1).

### SpoIVA polymers align directionally along the forespore surface

Most sporangia were sectioned along a median plane intersecting both forespore poles, allowing clear visualization of polar structures. In these tomograms, the most prominent SpoIVA objects were observed at the forespore poles and appeared as projections of linear polymers oriented perpendicular to the imaging plane (Fig. 2Ai-ii). Orthogonal views further confirmed their presence as linear densities aligned along the same direction within the depth of the section (Fig. 2Aii-iii), indicating that SpoIVA filament-like structures stripe the forespore poles. Interestingly, at the tip of the mother-proximal pole, some polymers appeared to merge or branch (Fig. 2Aii-iii), suggesting that these regions may be rich in nucleation spots.

**Figure 2.**
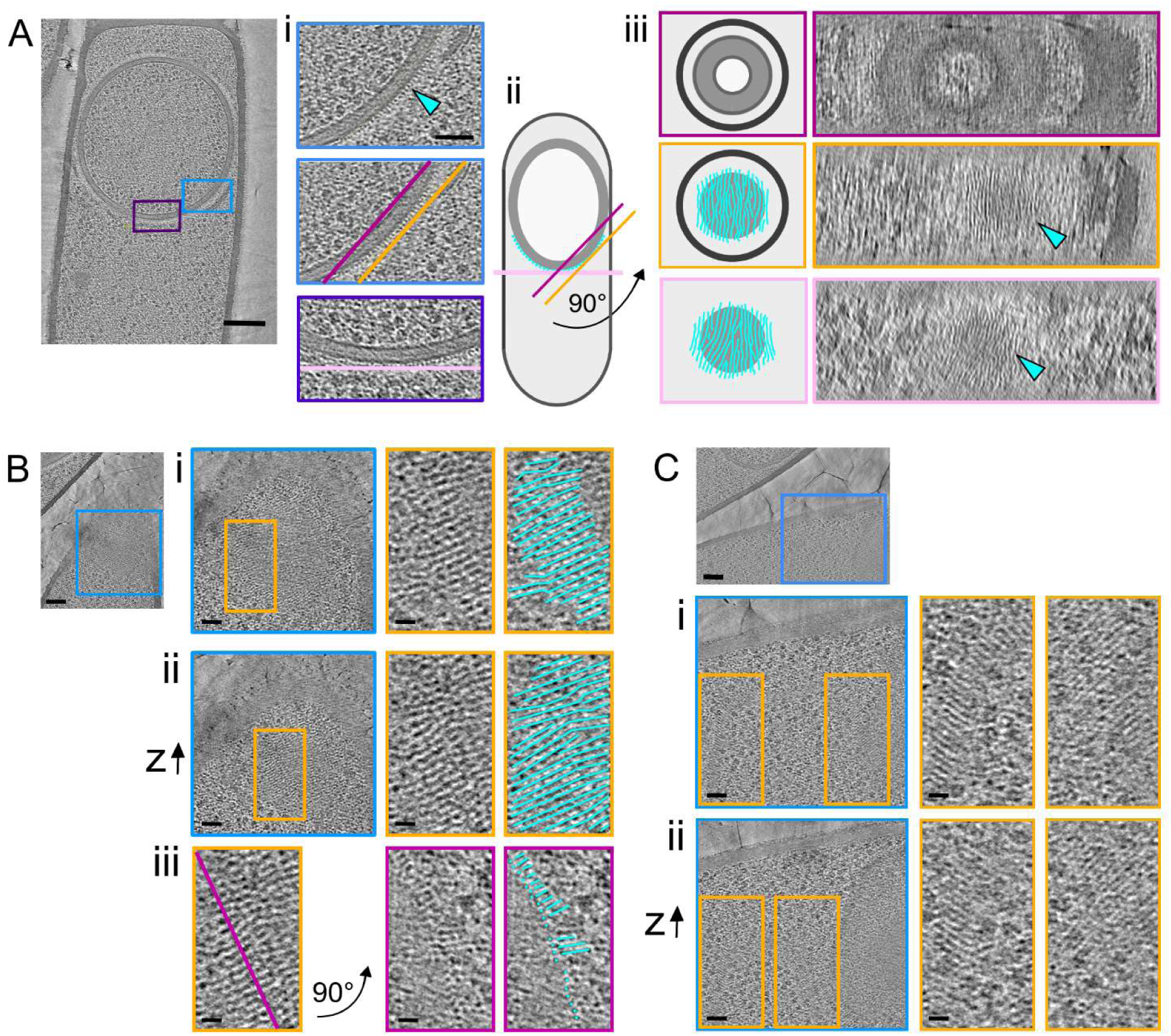
Spatial organization of SpoIVA tracks around the forespore. **A-C.** Slices through cryo-electron tomograms of *B. subtilis* sporangia, presented as full views (scale bars, 200 nm) or magnified views of the forespore surface (scale bars, 20 nm). Insets highlight the enlarged regions, and lines indicate the orientation of orthogonal planes where applicable. **A.** Analysis of the 3D organization of the SpoIVA layer in a forespore sectioned along its median plane. SpoIVA polymers (cyan arrowheads) appear as crenelated structures (**i-ii**) when observed in planes parallel to the section, or as filaments when viewed in planes orthogonal to the section (**ii**-**iii**). Panel (**ii**) shows a schematic illustrating the visible SpoIVA particles around the forespore and the various observation planes, with lines color-coded as in panels (**i**) and (**iii**). In panel (**iii**), the mother cell and forespore envelopes are shown as black and grey circles or disks, respectively. **B-C.** Slices through cryo-electron tomograms of forespores sectioned near their surface, showing SpoIVA polymers (highlighted by cyan lines). Panels (**i**) and (**ii**) provide magnified views of the forespore surface at incremental depths and yellow insets zoom in on SpoIVA tracks (scale bars, 50 nm. In panel (**iii**), polymers striping the forespore surface are observed in a plane orthogonal to the section (magenta line).

To examine the orientation of the SpoIVA polymers in the forespore equatorial region, we took advantage of 4 medium-magnification tomograms in which forespores were sectioned near their surface (Figs. 2B-C). In these tomograms, filament-like structures ran predominantly parallel to one another (Fig. 2Bi-ii, 2Ci-ii). Orthogonal views revealed regularly spaced, point-like densities (4.5 ± 0.6 nm in diameter, n = 14) (Fig. 2Biii), consistent with the dimensions of the SpoIVA objects described above. Beyond this uniform arrangement, in two tomograms, the orientation of linear polymers relative to the sporangium longitudinal axis varied across distinct regions of the forespore surface (Figs. 2C).

Collectively, these observations suggest that SpoIVA polymers nucleate within the mother-cell proximal polar zone and extend directionally along the forespore surface toward the distal pole, with local variations in filament orientation in the equatorial region.

### SpoIVA forms dimers through association of its stalk and base regions

To investigate SpoIVA organization within linear polymers, we generated monomeric and oligomeric AlphaFold3 models (Figs. 3, S2 and S3) (Abramson et al., 2024). The overall shape of the monomeric SpoIVA model (mean pLDDT (per-atom) confidence score of 83.03%) is reminiscent of a statue on a pedestal (Fig. S3A-B). This structure, with a 12-nm height, includes the N-terminal ATPase domain (∼ 5-nm radius, similar to the globular region of the averaged SpoIVA objects), connected to a stalk oriented perpendicular to a C-terminal base region (Figs. 3A and S3A).

**Figure 3.**
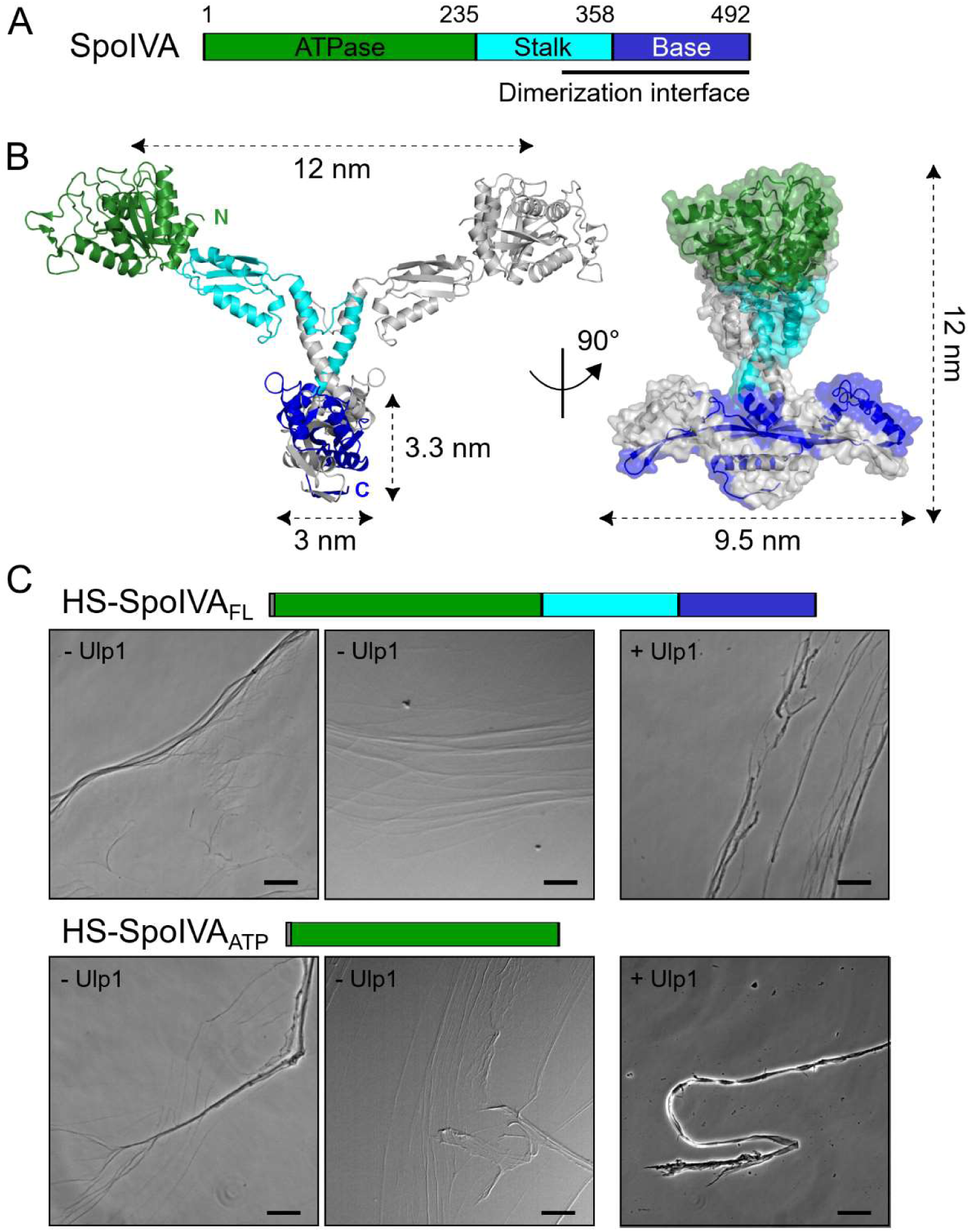
SpoIVA dimerization and filament formation. **A.** Domain organization of SpoIVA, with residue numbering based on the *B. subtilis* 168 sequence. The the AlphaFold3-predicted structure of the SpoIVA monomer, highlighting the N-terminal globular ATPase domain (green), the stalk region (cyan), and the base region (blue). **B.** Ribbon diagrams and surface representation (right panel) showing orthogonal views of the AlphaFold3-predicted SpoIVA dimer. One protomer harbors the color code of panel **A**, while the second protomer is shown in grey. Domain dimensions and the positions of the N- and C-termini are annotated. **C.** Phase-contrast images of purified His_6_-SUMO-SpoIVA_FL_ (HS-SpoIVA_FL_, residues E2-L492) and His_6_-SUMO-SpoIVA_ATP_ (HS-SpoIVA_ATP_, residues E2-E235) after 24-h incubation at 37 °C in the presence of 4 mM ATP, with or without HS-tag cleavage by the Ulp1 protease. Scale bars, 200 nm.

Consistent with *in vitro* dimerization reports (Updegrove et al., 2021), AlphaFold3 predictions for even and odd SpoIVA multimers consistently converged toward a high-confidence dimer (pLDDT ∼ 80%) (Fig. 3B and S3C). Interestingly, the ATPase domains, positioned about 12 nm apart, are not involved in dimerization. Instead, AlphaFold3 predicts extensive hydrophobic and electrostatic interactions between the stalk and base regions, with a buried surface of about 6,000 Å ² (Fig. S4). The stalks interact through helices α8, α11 and α12, forming a composite six-helix bundle (Figs. S3D and S4A-B). The tightest interactions are established by strands β17 and β18 of the base regions, whose association across the two protomers results in a composite antiparallel β-sheet, around which all helices of the base region also contribute to the dimeric interface (Fig. S3D and S4C). Finally, a small, composite antiparallel β-sheet is formed by strands β21 in the lower base region (Fig. 3B, S3D and S4D-E). This network forms a dimeric base of about 3.3 nm in height, 3.0 nm in width, and 9.5 nm in length.

To test the AlphaFold3 dimer model, we purified His_6_-SUMO (HS)-tagged recombinant constructs of SpoIVA, including the full-length protein (SpoIVA_FL_, encompassing residues E2 to L492), the ATPase domain alone (SpoIVA_ATP_, encompassing residues E2 to E235), and two full-length variants harboring mutations in conserved interface residues located in the stalk (SpoIVA_FL_(L337E-M340E)) or base (SpoIVA_FL_(L377E-I403E)) regions (Fig. S5A). Following cleavage of the HS-tag, mass photometry revealed dimers (100 ± 1 kDa) as the primary species (theoretical MW of 110 kDa), with a minor population (187 ± 7 kDa) potentially corresponding to SpoIVA tetramers (Fig. S5B). Size-exclusion chromatography coupled to multi-angle laser light scattering (SEC-MALLS) of SpoIVA_FL_ confirmed a dominant, gaussian-shaped elution peak at ∼ 11 mL with a flat molecular weight (MW) trace of 91.0 ± 13.5 kDa, consistent with a SpoIVA homodimer (Fig. S5C). A high-MW species, eluting at 7 mL, might correspond to polymerized or aggregated SpoIVA. By contrast, SEC-MALLS of the ATPase domain showed a single species with an apparent MW of 25.4 ± 4.0 kDa (Fig. S5D), matching the theoretical MW of a monomer (26 kDa), and confirming that the ATPase domain alone does not mediate dimerization. Dimerization was destabilized in both the stalk and base mutants, with SpoIVA_FL_(L337E-M340E) predominantly monomeric (apparent MW of 46 ± 1 kDa), and SpoIVA_FL_(L377E-I403E) exhibiting a mix of dimers (apparent MW of 82 ± 1 kDa) and monomers (apparent MW of 43 ± 1 kDa) (Fig. S5E-F). Altogether, these results indicate that SpoIVA dimerizes via its stalk and base regions.

### SpoIVA polymerization does not require dimerization

To determine whether SpoIVA polymerization requires pre-assembly into dimers, we examined the ability of His_6_-SUMO-SpoIVA (HS-SpoIVA) variants to form filaments *in vitro*. These experiments could not be performed with proteins pre-cleaved from the HS tag, as they aggregated in the polymerization buffer even in the absence of ATP. Instead, polymerization assays were conducted using HS-tagged proteins, either with or without the Ulp1 protease, which cleaves the SUMO tag. Under all conditions tested and in the presence of ATP, both HS-SpoIVA_FL_ and HS-SpoIVA_ATP_ formed filaments and filament bundles (Fig. 3C). However, filaments formed by the ATPase domain were less abundant when Ulp1 was added, likely because tag removal reduces protein solubility and promotes aggregation. Notably, similar filaments and filament bundles were obtained with a previously reported His_6_-Thr-SpoIVA construct (Fig. S6A; Ramamurthi and Losick, 2008), showing that the SUMO tag does not influence SpoIVA polymerization.

Consistent with the polymerization capability of HS-SpoIVA_ATP_, the interface mutants HS-SpoIVA_FL_(L337E-M340E) and HS-SpoIVA_FL_(L377E-I403E) also formed filaments, although these were considerably sparser, particularly in the presence of Ulp1 (Fig. S6B-C). These constructs exhibit a tendency to aggregate, indicating that proper dimerization contributes to SpoIVA solubility *in vitro*. Together, these results show that the ATPase domain alone is sufficient to drive filament formation, whereas dimerization via the stalk and base regions harbors another function.

### Dimerization is critical for SpoIVA localization

To assess the physiological relevance of SpoIVA dimerization, we built *B. subtilis* strains expressing the *spoIVA(L337E-M340E)* and *spoIVA(L377E-I403E)* variants under the control of the *spoIVA* promoter. Both mutant strains phenocopied Δ*spoIVA* in heat-kill assays, failing to produce heat-resistant spores. Since both SpoIVA variants were expressed and stable (Fig. S7), we conclude that SpoIVA dimerization plays a critical role in spore development.

To determine whether dimerization influences SpoIVA localization, we examined the fluorescence pattern of a previously characterized mYPET-SpoIVA fusion in strains expressing the *spoIVA* variants (Luhur et al., 2020). As the mYPET-SpoIVA fusion is not fully functional (Luhur et al., 2020), these experiments were performed in merodiploid strains containing the mYPET-SpoIVA fusion and untagged SpoIVA variants. In the otherwise wild-type background, mYPET-SpoIVA localized to the engulfing membrane at T2.5 (2.5 h after entry into sporulation) (Fig. 4A), with stronger fluorescence in the mother-cell proximal region of engulfed forespores. By T3.5, the mYPET-SpoIVA signal became more uniformly distributed across the forespore surface (Figs. 4B).

**Figure 4.**
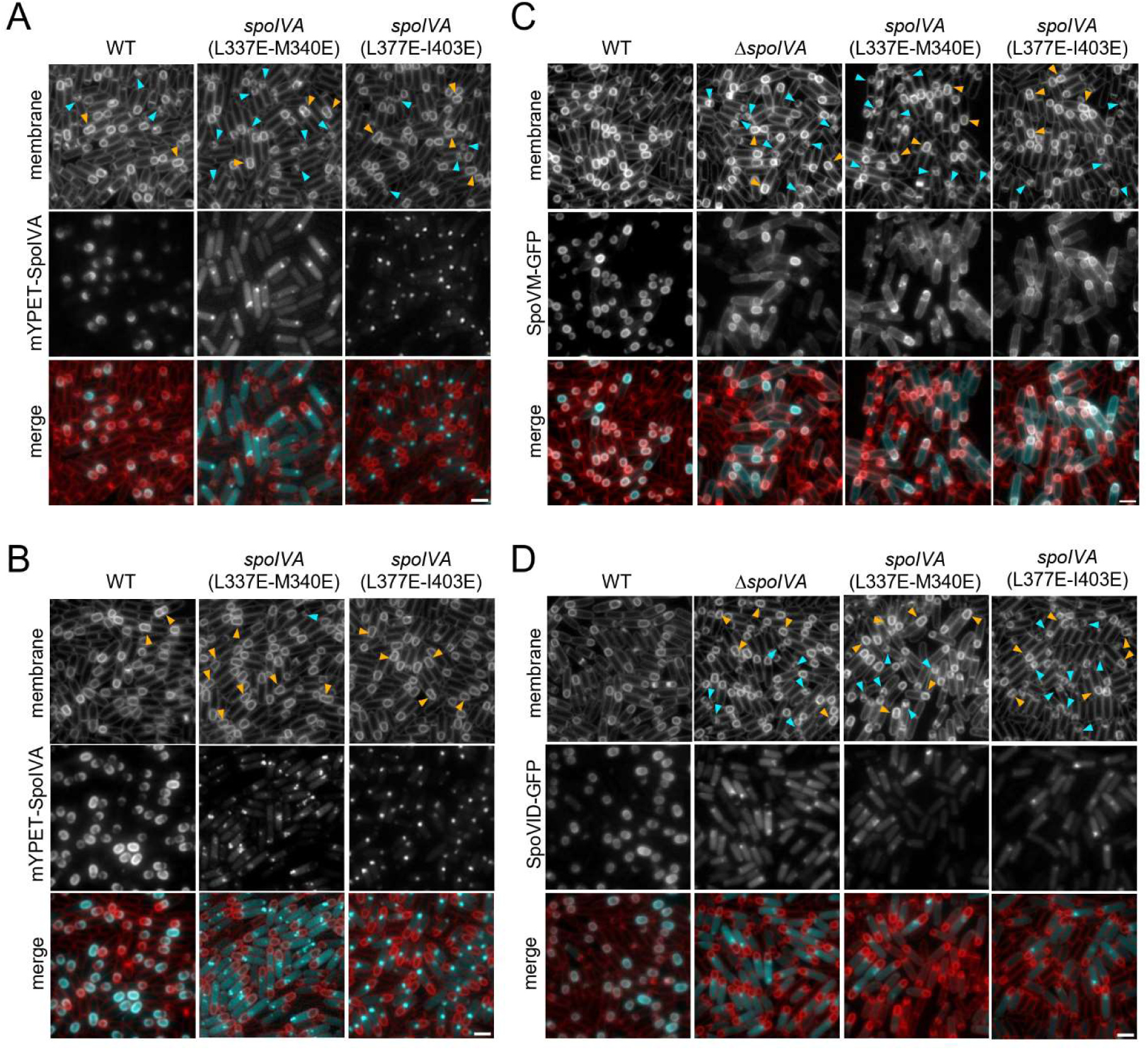
Disruption of SpoIVA dimerization alters its localization and function. **A-B.** Fluorescence microscopy showing the localization of mYPET-SpoIVA (left panels), mYPET-SpoIVA(L337E-M340E) (middle panels), or mYPET-SpoIVA(L377E-I403E) (right panels) in wild-type (WT), *spoIVA(L337E-M340E)*, or *spoIVA(L377E-I403E)* backgrounds, respectively. **C-D.** Fluorescence microscopy showing the localization of SpoVM-GFP (**C**) or SpoVID-GFP (**D**) in WT, *spoIVA(L337E-M340E)*, or *spoIVA(L377E-I403E)* backgrounds. Images were acquired 2.5 h (**A**, **C**, **D**) or 3.5 h (**B**) after the onset of sporulation. The mYPET and GFP signals are false-coloured cyan in the merged images. Cell membranes were visualized with the TMA-DPH fluorescent dye, and are false-coloured red in the merged images. Septal membrane bulges are highlighted with blue triangles, and indented cells are highlighted with orange triangles. Scale bars, 2 µm.

In contrast, the mYPET-SpoIVA(L337E-M340E) stalk mutant failed to localize around the forespore, instead forming clusters near the forespore surface or in the mother-cell cytoplasm, and harboring diffuse localization in the mother-cell cytoplasm (Fig. 4A-B). The mYPET-SpoIVA(L377E-I403E) base mutant primarily mislocalized as single foci at the mother-cell proximal side of the forespore membrane, with minimal cytoplasmic signal (Fig. 4A-B). Mutations disrupting the dimerization interface in either the stalk or the base region thus alters SpoIVA localization around the forespore.

### Dimerization is critical for SpoIVA function

We next investigated how SpoIVA dimerization influences the localization of two direct partners, SpoVM and SpoVID, using previously described GFP fusions (Luhur et al., 2020). Consistent with earlier observations (Ramamurthi et al., 2006; Wang et al., 2009), SpoVM-GFP delineated the engulfing membrane in wild-type sporangia but was partially mislocalized to the mother-cell membrane in the absence of SpoIVA (Figs. 4C and S8A). A similar mislocalization pattern was observed in both dimerization interface mutants, with a more severe defect in the *spoIVA(L377E-I403E)* background. These results indicate that disruption of SpoIVA dimerization, particularly within the base region, compromises SpoVM localization.

In wild-type sporangia, SpoVID-GFP localized around the forespore at T2.5 and T3.5. By contrast, in Δ*spoIVA*, *spoIVA(L337E-M340E*) and *spoIVA(L377E-I403E*) strains, SpoVID-GFP exhibited diffuse fluorescence signal throughout the mother cell cytoplasm, together with discrete foci near the mother cell-forespore interface (Figs. 4D and S8B). Given that coat layers form swirls in the mother cell cytoplasm of Δ*spoIVA* cells (Bauda et al., 2024; Catalano et al., 2001), the SpoVID-GFP foci likely correspond to coat material detached from the forespore surface. Together, these findings indicate that SpoIVA dimerization is required for proper propagation of the basement layer and attachment of the coat around the forespore.

Further supporting the importance of SpoIVA dimerization for spore development, fluorescence microscopy analysis of *spoIVA(L337E-M340E)* and *spoIVA(L377E-I403E)* mutants revealed membrane bulges and asymmetric positioning of the engulfing membrane front (Figs. 4 and S8). This phenotype is also observed in Δ*spoIVA* sporangia (Fekade et al.) and was previously described in mutants lacking the DMP complex, which is essential for septum remodeling and engulfment (Khanna et al., 2019; Morlot et al., 2010). In addition, consistent with what has been reported for Δ*spoIVA* (Fekade et al.), all *spoIVA* variants exhibited forespore indentation in a subset of cells, after engulfment completion (Figs. 4 and S8). To examine the cell envelope architecture of *spoIVA* mutant sporangia, we performed cryo-ET and TEM analysis at T2 and T3. Both partially and fully engulfed Δ*spoIVA* forespores were sandwiched between mother-cell envelope protrusions that were located in the mother cell-distal quarter of the sporangium (Fig. S9), thus consistent with the position of the asymmetric septum formed at the onset of sporulation. Tomograms of engulfed Δ*spoIVA* forespores showed slight compression by these envelope outgrowths (Fig. S9B), aligning with the indentation phenotype observed by fluorescence microscopy. TEM of resin-embedded sections from *spoIVA* dimerization mutants confirmed the presence of both membrane bulges, characteristic of incomplete septum degradation (Khanna et al., 2019; Morlot et al., 2010), and the forespore indentation phenotype (Fig. S10).

Together, these findings suggest that forespore bulging and indentation, which are observed in all *spoIVA* variants, stem from impaired septal peptidoglycan degradation, implicating SpoIVA dimerization in the engulfment process.

## Discussion

In this study, cellular cryo-ET analysis of *B. subtilis* spores reveals that SpoIVA assembles into oriented, uniformly spaced linear polymers embedded in the OFM, likely in complex with SpoVM. The striking linearity and regularity of these structures, combined with *in vitro* reconstitution demonstrating their static nature (as shown here and in (Ramamurthi and Losick, 2008)), strongly suggest that they function as “tracks” rather than “filaments”, the latter typically associated with dynamic structures.

The temporal emergence of SpoIVA assemblies at the onset of engulfment (Bauda et al., 2024) indicates that oligomer nucleation coincides with the initiation of septal peptidoglycan thinning. Supporting this, we observed branching tracks at the forespore pole, likely representing nucleation sites within the asymmetric septum. Furthermore, the consistent perpendicular orientation of polar tracks in median-plane tomograms aligns with the direction of the engulfing membrane. Collectively, these observations suggest that SpoIVA tracks polymerize from confined nucleation sites located at the mother cell-proximal forespore pole and extend toward the distal pole following the direction of engulfment (Fig. 5).

**Figure 5.**
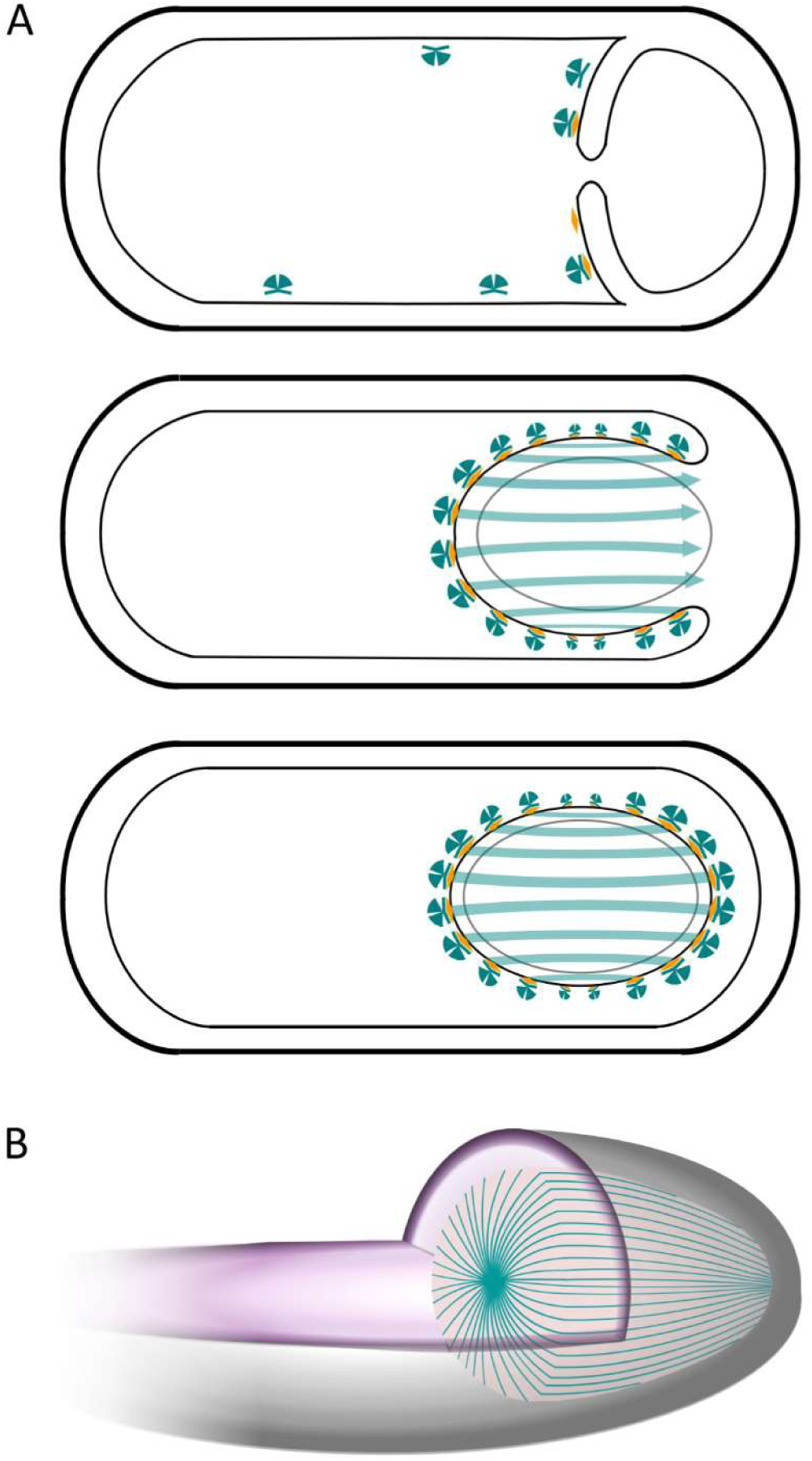
Proposed model for the polymerization and organization of SpoIVA tracks. **A.** Schematics of the SpoIVA oligomerization mechanism. During early engulfment, SpoIVA dimers (cyan) sample the membrane until being stabilized in the OFM through interaction with SpoVM (yellow). Since SpoVM preferentially associates with positively curved membranes, SpoIVA-SpoVM complexes accumulate at the mother-cell proximal forespore pole. As engulfment progresses, SpoIVA polymers extend toward the mother-cell distal pole. ATP binding and hydrolysis promote and stabilize track formation. **B.** Three-dimensional illustration of an engulfed forespore within the mother cell. The SpoIVA tracks (cyan lines) radiate from the mother-cell proximal pole and stripe the forespore surface.

AlphaFold3 modeling of SpoIVA identified three distinct regions: a globular N-terminal ATPase domain, an elongated stalk, and a large C-terminal base. Although the resolution of our track reconstruction did not permit tracing the protein backbone, the predicted SpoIVA architecture aligns well with the experimental polymer dimensions. The base, which interacts with SpoVM (Ramamurthi et al., 2006), would form the region embedded in the OFM, positioning the globular ATPase domain (predicted radius ∼ 5 nm) about 10 nm above the OFM. These predictions are consistent with the dimensions (∼ 4.2 nm in height and 5.0 nm in width) and position (∼ 9 nm above the OFM) of the globular region of the averaged structures. However, the length of the OFM-embedded region estimated from cryo-ET (∼ 4.4 nm) is roughly half the longest dimension of the base region predicted by AlphaFold3 (∼ 9.5 nm). This discrepancy suggests that the long axis of the basal region is not perpendicular to the polymerization axis, implying a tilted arrangement of SpoIVA dimers within the tracks.

SpoVM is known to distribute SpoIVA around the OFM (Ramamurthi et al., 2006), yet SpoIVA retains partial OFM localization in the absence of SpoVM, indicating an intrinsic affinity for hydrophobic and positively curved surfaces (Ramamurthi et al., 2006; Wu et al., 2015). The dimeric AlphaFold3 model of SpoIVA, supported by biophysical and functional assays, provides a plausible explanation: the two C-termini form a β-sheet (β21-β21) with a hydrophobic exposed face, potentially acting as a membrane-interacting appendage. This might allow SpoIVA dimers to localize to the OFM, where they are further stabilized by interaction with SpoVM (Peluso et al., 2019). Consistent with this, disruption of the dimerization interface abolishes SpoIVA ability to delineate the OFM, instead forming foci at the mother cell-forespore interface, as reported in Δ*spoVM* sporangia (Ramamurthi et al., 2006). Conversely, SpoVM mislocalizes in Δ*spoIVA* and *spoIVA* dimerization mutants, demonstrating that SpoIVA dimerization is as critical for SpoVM localization as the presence of the protein itself. SpoIVA dimerization mutants also exhibit SpoVID mislocalization, mimicking the phenotype of Δ*spoIVA* cells. Thus, SpoIVA dimerization is essential for basement layer assembly, encasement and subsequent coat formation.

The formation of stable SpoIVA dimers, however, does not appear crucial for polymerization, as the ATPase domain alone can polymerize *in vitro*. The reduced number of polymers observed with dimerization mutants likely stems from a tendency to aggregate rather than from a direct contribution of the stalk and base regions to filament formation. Instead, these regions may modulate the organization of the linear tracks. For instance, the width of the base region could enforce the uniform spacing between SpoIVA tracks. Since regular spacing persists throughout the polymer height (from the base to the ATPase domain), another component likely stabilizes the straight alignment of the tracks. The weak density observed between consecutive tracks suggests the presence of flexible molecules, such as glycans or intrinsically disordered protein regions, that likely occupy the inter-track space. SpoVID, with its long disordered region and C-terminal LysM domain required for SpoIVA interaction (Müllerová et al., 2009; Wang et al., 2009), emerges as a strong candidate. Its flexible region may intercalate between SpoIVA oligomers to maintain regular spacing. This hypothesis is consistent with the predicted localization of SpoVID’s LysM domain, which sequesters Lipid II to prevent premature cortex synthesis and, upon basement layer maturation, interacts with polymerized SpoIVA to release Lipid II (Delerue et al., 2022). Given that Lipid II is anchored in the OFM, this model positions the LysM domain near SpoIVA, with the disordered region potentially filling the space between adjacent tracks. Other OFM proteins, such as the DMP or the A-Q complex, may also contribute to track spacing.

Attempts to model SpoIVA oligomers using AlphaFold3 have not yet yield models consistent with the architecture of the *in-situ* tracks. Given the large interface involved in SpoIVA dimers, this assembly is likely very stable. In addition, *spoIVA* dimerization mutants exhibit more pronounced SpoVID localization defects than a *spoIVA(K30A)* catalytic mutant (Fekade et al.). This suggests that SpoIVA dimerization precedes polymerization, and that SpoIVA dimers constitute the forming unit of the linear polymers. We therefore propose a dual oligomerization mechanism, in which SpoIVA dimers anchor to the OFM via SpoVM, polymerize into linear tracks, and are stabilized by ATP hydrolysis, forming a scaffold for coat deposition.

Importantly, this work also shows that SpoIVA deletion or disruption of its dimerization leads to the expansion of forespore membranes into the mother-cell compartment and uneven progression of the engulfing membrane front. Such defects were initially described in mutants lacking the DMP complex, which drives engulfment by degrading septal PG at the leading edge of the engulfing membrane (Abanes-De Mello et al., 2002; Chastanet and Losick, 2007; Khanna et al., 2019; Morlot et al., 2010). Strikingly, Δ*spoIVA* and dimerization-defective mutants also display indentations of the mother-cell envelope toward the forespore, whereas in wild-type sporangia, similar protrusions are restricted to one side of the cell (Bauda et al., 2024). This indicates that SpoIVA limits the formation of these protrusions. In a parallel study, Fekade et al. also observed membrane bulges, asymmetric engulfment and forespore indentation in the Δ*spoIVA* and *spoIVA(K30A)* mutants, linking SpoIVA polymerization to septal peptidoglycan hydrolysis and engulfment (Fekade et al.). Together, these findings establish a functional interplay between SpoIVA dimers and tracks, septum hydrolysis and membrane dynamics during engulfment.

While further studies are required to dissect the molecular basis of SpoIVA oligomerization and the role of protein partners in the polymer organization, this work unravels the *in situ* architecture of the SpoIVA layer, providing the first direct evidence that SpoIVA assembles into linear tracks on the forespore surface. Beyond its established role in coat assembly, this work also shows that SpoIVA oligomerization is critical for septum remodeling and engulfment, revealing a previously underappreciated structural-function relationship for this universally conserved coat protein.

## Materials and Methods

### Plasmids, bacterial strains and culture conditions

Strains and plasmids used in this study are listed in Extended Table 1. All *B. subtilis* strains were derived from the auxotrophic strain 168 (Zeigler et al., 2008). Cryo-electron tomography data were acquired on a Δ*spoIVB* strain, as in (Bauda et al., 2024). *B. subtilis* cells were grown in CH medium at 37 °C to OD_600nm_ ∼ 0.3-0.5. Sporulation was then induced by resuspension in sporulation medium at 37°C, according the method of Sterlini-Mandelstam (Harwood and Cutting, 1990).

### Cryo-FIB/SEM milling

Cryo-FIB milling was performed on plunge frozen *B. subtilis* cells according to the protocol previously described (Bauda et al., 2024). FIB micro-machining of lamellae in cryogenic conditions was performed on a Zeiss Crossbeam 550A equipped with a Quorum cryostation. In brief, 150 to 300 nm-thick lamellae (10 × 10 µm) were made using rectangular milling patterns and beam current of 1 nA to 0.37 nA for rough milling, 100 pA to 50 pA for fine milling and 30 pA for polishing. On each side of the lamella, 20 x 0.2 µm stress-relief trenches were prepared with a current of 1.5 nA. Detailed cryo-FIB/SEM workflow for bacterial cell section is described in (Bauda et al., 2024).

### Cryo-electron tomography

Tilt series were acquired on a 300 kV TITAN KRIOS electron microscope (TFS) equipped with a post-column energy filter and K3 direct detection camera (Ametek) for medium-magnification tomograms at the ESRF CM01 beamline (Grenoble, France) (Kandiah et al., 2019). For high-magnification tomograms used for subtomogram averaging (STA), tilt series were acquired on a 300 kV TITAN KRIOS electron microscope (Thermofisher) equipped with a post-column energy filter and K2 direct detection camera (Ametek) at the CEITEC (Brno, Czech Republic). The specimen were tilted from - 60 to + 60 ° with an increment of 3.0 ° (medium-magnification datasets), or from - 50 to + 50 ° with an increment of 2.5 ° (high-magnification datasets) based on the dose symmetric Hagen scheme. The 0 ° of the tilt series was defined considering the intrinsic tilt of the lamellae caused by the milling angle. The tilt series were acquired using SerialEM(Mastronarde, 2005) or Tomo5 (TFS). The images were recorded at a defocus of - 10 µm and nominal magnification of 19,500 (pixel size of 4.5 Å) on the K3 camera, or at a defocus of - 1.5 or - 4 μm at nominal magnifications of 81,000 (pixel size of 1.8 Å) on the K2 camera, with a cumulative dose of ∼ 80-90 e-/Å _2_.

### Image Processing and segmentation

Tilt series were first drift-corrected using MotionCor2 (Zheng et al., 2017). Alignment of the tilt series was then performed in the IMOD software package using the patch tracking method (Mastronarde, 2006). Tomograms were reconstructed in IMOD, using weighted back projection. For STA, PEET was employed (Nicastro et al., 2006). SpoIVA particles were visually identified in individual tomograms, and their orientations relative to the OFM were defined using 3dmod to generate model points, the initial motive list, and particle rotation axes (Mastronarde, 2006). A total of 1,051 particles were averaged using a box size of 45×45×45 pixels and a pixel size of 6.9 Å. The 3D rendering of the averaged structures was performed in ChimeraX (Goddard et al., 2018). Finally, the dimensions of objects present in the tomograms were manually measured in 3dmod (Mastronarde, 2006).

### Transmission electron microscopy

Ten mL of cell culture was collected at desired time points and pelleted at 13,000 rpm for 5 min, after which they were transferred into primary fixative (4% glutaraldehyde in 0.1 M sodium cacodylate buffer) and incubated overnight at 4 °C. Samples were then stained with 1% osmium tetroxide for 1 h, followed by washing. After stepwise dehydration in 25%, 50%, 75% and 100% acetone, they were infiltrated with 50% resin for 1 h, followed by 100% resin for 24 hours. The Agar LV resin was cured at 60 °C degrees overnight. After ultrathin sectioning on an RMC ultramicrotome sections were post-stained in 2% uranyl acetate for 2 min and 1.5% lead citrate for 1 min. Samples were imaged in a JEOL JEM2100Plus with Gatan OneView CMOS camera.

### Production and purification of SpoIVA recombinant constructs

The previously reported His_6_-Thr-tagged, full-length SpoIVA recombinant construct (residues M1 to L492, HT-SpoIVA_FL_) (Ramamurthi and Losick, 2008), along with all His_6_-SUMO-tagged versions of SpoIVA (Extended Table 1) were overexpressed in *E. coli* BL21 (RIL). Cells were grown in 2 L of Terrific Broth (TB) medium supplemented with 50 μg⋅mL^−1^ kanamycin at 37 °C until reaching an OD_600nm_ of 1.5. Protein expression was induced overnight at 25°C by adding 0.5 mM IPTG (isopropyl β-D-1-thiogalactopyranoside). Cells were harvested by centrifugation, resuspended in 1/25^th^ volume of buffer A [50 mM Tris-HCl pH 8.0, 500 mM NaCl, 25 mM imidazole, 1 mM TCEP, 10% (vol/vol) glycerol] supplemented with the Complete protease inhibitor cocktail (Roche), and lysed via six passages through a Microfluidizer M110-P (Microfluidics) at 20,000 psi. Cell debris were removed by centrifugation at 40,000 × *g* for 30 min at 4°C. HT and HS fusions were purified using 8.0 mL of Ni-NTA agarose resin (Qiagen) equilibrated in buffer A. After sample loading and extensive washing with buffer A, tagged proteins were eluted over 10 column volumes using a linear 50–500 mM imidazole gradient in buffer B [50 mM Tris-HCl pH 8.0, 300 mM NaCl, 1 mM TCEP, 10% (vol/vol) glycerol]. The HT-SpoIVA_M1-L492 protein and half of the HS-tagged samples were aliquoted and flash frozen with liquid nitrogen for polymerization assays. To remove the HS-tag, samples were dialyzed overnight at 4°C in buffer B without imidazole and 1:250 molar ratio of His_6_-Ulp1 protease (Uehara et al., 2010). Cleavage reactions were passed through Ni-NTA resin to remove free HS-tags and His_6_-Ulp1. Flow-through fractions containing untagged SpoIVA were concentrated using Amicon ultra centrifugal filter units (10-kDa molecular weight cutoff (MWCO), Millipore), and further purified via size-exclusion chromatography (SEC) on an ENrich™ SEC 650 10 x 300 column (Bio-Rad) equilibrated in buffer C [25 mM Tris-HCl pH 8.0, 150 mM NaCl, 1 mM TCEP]. Purified proteins were concentrated again using 10-kDA MWCO Amicon ultra centrifugal filter units.

### SEC-MALLS analyses

SEC coupled with multi-angle laser light scattering (MALLS), refractometry, and UV-280 nm detection was used to determine the apparent molecular mass of SpoIVA in solution. SEC was carried out at 20 °C using a Superdex 200 10/300 column (Cytiva) equilibrated in 25 mM Tris-HCl pH 8.0 and 150 mM NaCl. A volume of 50 µL of protein solution at 1-8 mg⋅mL^−1^ was injected at a constant flow rate of 0.5 mL⋅min^1^. MALLS and differential refractive index detection were carried out using a MiniDAWN-TREOS detector (Wyatt Technology Corp.) with a 690 nm laser and an Optilab T-rEX detector (Wyatt Technology Corp.), and an OMNISEC system (MALVERN) equipped with right-angle light scattering (RALS), low-angle light scattering (LALS), viscometer, and refractive index detectors. Weight-averaged molecular mass was determined using ASTRA software (Wyatt Technology Corp.) or OMNISEC software version v11.40 (MALVERN).

### Mass photometry

The landing of SpoIVA on high-precision glass coverslips (24 × 50 mm^2^, No. 1.5H; Marienfeld) was recorded using a Refeyn OneMP mass photometry system (Refeyn Ltd.). Coverslips were cleaned according to the manufacturer’s protocol (Refeyn Ltd.). To maintain the shape of the sample droplet, reusable 6-well self-adhesive silicone gaskets (Refeyn Ltd.) were used. Movies were acquired for 60 sec (6,000 frames) using AcquireMP software version v2.4.2 (Refeyn Ltd.). Contrast-to-mass calibration was performed using a mixture of standard proteins with molecular weights of 66, 146, 480 and 1,048 kDa (Native Marker, Invitrogen). Data were analyzed using DiscoverMP software version v2.4.2 (Refeyn Ltd.), with a binning of 5 frames. The mean peak contrast per molecular weight was determined by Gaussian fitting. Sample droplets were prepared by adding 1 µL of protein solution to 19 µL of analysis buffer, resulting in a final droplet volume of 20 µL and a protein concentration of 13.5 nM. The experimental mass of SpoIVA proteins was derived from the calibration curve by averaging three independent measurements.

### *In vitro* SpoIVA polymerization assays

Before polymerization assays, purified HT-SpoIVA and HS-SpoIVA proteins were dialyzed against buffer D [50 mM Tris·HCl pH 7.5, 150 mM NaCl, 1 mM DTT] overnight at 4 °C. 15 µM of HT-SpoIVA or HS-SpoIVA samples were then incubated in a 55-μL reaction volume in buffer C [50 mM Tris·HCl pH 7.5, 400 mM NaCl, 1 mM DTT, 1 mM MgCl_2_] in the presence or absence of 4 mM ATP at 37 °C for 24 h. Samples were centrifuged at 20,000 × *g* for 2 min at 20 °C. For optical microscopy analysis, 5 µL of polymerized material were removed from the bottom of the tube, loaded onto a clean glass microscope slide, and immobilized with a clean glass coverslip. A 2-kg weight was applied onto the coverslip for 15 min to allows SpoIVA polymers to settle on the microscopy slide. Samples were imaged using a motorized two-deck Olympus IX83 optical microscope equipped with a UPFLN 100X O-2PH/1.3 objective and an ORCA-Flash4.0 Digital sCMOS camera from Hamamatsu. Images were acquired using the Volocity software package.

### Fluorescence microscopy

Live-cell fluorescence imaging was conducted by placing cells on a 2% (w/v) agarose pad made with resuspension medium and mounting them on a Gene Frame (Thermofisher). Specifically, 250 µL of culture was collected at a defined time point, pelleted by centrifugation, and resuspended in 10 µL of resuspension medium containing 0.05 mM TMA-DPH [1-(4-trimethylammoniumphenyl)-6-phenyl-1,3,5-hexatriene p-toluenesulfonate]. A 2-µL aliquot of the cell suspension was then spread onto the agarose pad, and a coverslip was placed on the adhesive Gene Frame.

Cells were imaged using a Zeiss Axio Observer 7 microscope equipped with a Plan-Apochromat 100×/1.4 Oil Ph3 objective lens and a Colibri 7 Type R[G/Y]CBV-UV fluorescent light source. Images were captured with a Zeiss Axiocam 712 mono camera. The TMA-DPH membrane dye and GFP were excited using a Zeiss Axio 92HE filter with 100-ms exposure time. YFP and CFP fluorescence were acquired using a Zeiss Axio 108HE filter with 150-ms and 300-ms exposure, respectively.

### Immunoblot analysis

Whole-cell lysates from sporulating cells were prepared as previously described (Rodrigues et al., 2016). Samples were collected 2.5 h after the onset of sporulation and heated for 15 min at 50 °C prior to loading. Equivalent loading was based on OD600. Samples were separated on a 12.5% polyacrylamide gel and transferred to a PVDF membrane. Membranes were blocked in 5% non-fat milk with 0.5% Tween-20 for 1 h. Blocked membranes were probed with anti-SpoIVA (1:50000) primary antibody diluted into PBS with 0.05% Tween-20 containing 3% Bovine Serum Albumin (w/v) (Sigma) at 4 °C overnight. Primary antibodies were detected with horseradish-peroxidase anti-rabbit antibodies (BioRad) and detected with ECL Prime Western Blotting Detection reagent (Amersham) as described by the manufacturer.

## Data availability

The data that support this study are available from the corresponding authors upon request. The dataset that are necessary to interpret, verify and extend the research in the article are accessible through the Electron Microscopy Data Bank (EMDB) under accession codes EMD-XX, and through the Zenodo repository https://doi.org/XX.

## Acknowledgments

We thank members of the Morlot and Schoehn laboratories, and members of the electron microscopy and biophysical platforms, for support, advice and encouragement. We thank P.H. Jouneau for support and advices acknowin cryo-FIB/SEM milling. IBS acknowledges integration into the Interdisciplinary Research Institute of Grenoble (IRIG, CEA). This work used the platforms of the Grenoble Instruct-ERIC center (ISBG; UAR 3518 CNRS-CEA-UGA-EMBL) within the Grenoble Partnership for Structural Biology (PSB), supported by FRISBI (ANR-10-INBS-0005-02) and GRAL, financed within the University Grenoble Alpes graduate school (Ecoles Universitaires de Recherche) CBH-EUR-GS (ANR-17-EURE-0003). The IBS/ISBG electron microscope facility is supported by the Auvergne-Rhône-Alpes Region, the Fondation Recherche Médicale (FRM), the fonds FEDER and the GIS-Infrastructures en Biologie Santé et Agronomie (IBISA). We thank Guy Schoehn for the establishment of the EM facility. This work also benefited from access to beamline CM01 at the European Synchrotron Radiation Facility. We acknowledge cryo-electron microscopy and tomography core facility CEITEC MU of CIISB, Instruct-CZ Centre, supported by MEYS CR (LM2023042) and European Regional Development Fund-Project “UP CIISB” (No. CZ.02.1.01/0.0/0.0/18_046/0015974), Instruct-ERIC (PID1697 and PID17035), and iNEXT-Discovery (project number 871037) funded by the Horizon 2020 program of the European Commission. We acknowledge the Midlands Regional Cryo-EM Facility, hosted at the Warwick Advanced Bioimaging Research Technology Platform, for use of the JEOL 2100Plus, supported by MRC award reference MC_PC_17136. We acknowledge Danae Morales Angeles for construction of pDMÅ06. E.B. received funding from GRAL, a program from the Chemistry Biology Health (CBH) Graduate School of University Grenoble Alpes (ANR-17-EURE-0003), C.D.A.R. from the Biotechnology and Biological Sciences Research Council (grant BB/X008533/1) (https://www.ukri.org/councils/bbsrc) and B.F. and K.C. from the Midlands Integrative Biosciences Training Partnership (https://warwick.ac.uk/fac/cross_fac/mibtp/) (grant number BB/T00746X/1).

## Author Contributions

EB, CDAR and CM designed research; EB, BF, LB, BG, SD, KC, AL, GE, DF, CM, JN, GS and ChM performed experiments; EB, EN, CDAR and CM analyzed data, JM and GS provided support for FIB/SEM and electron microscope access and training; EB and CM wrote the manuscript; EB, CDAR and CM revised the manuscript.

## Extended Data Figures and Legends

**Extended data Figure 1.**
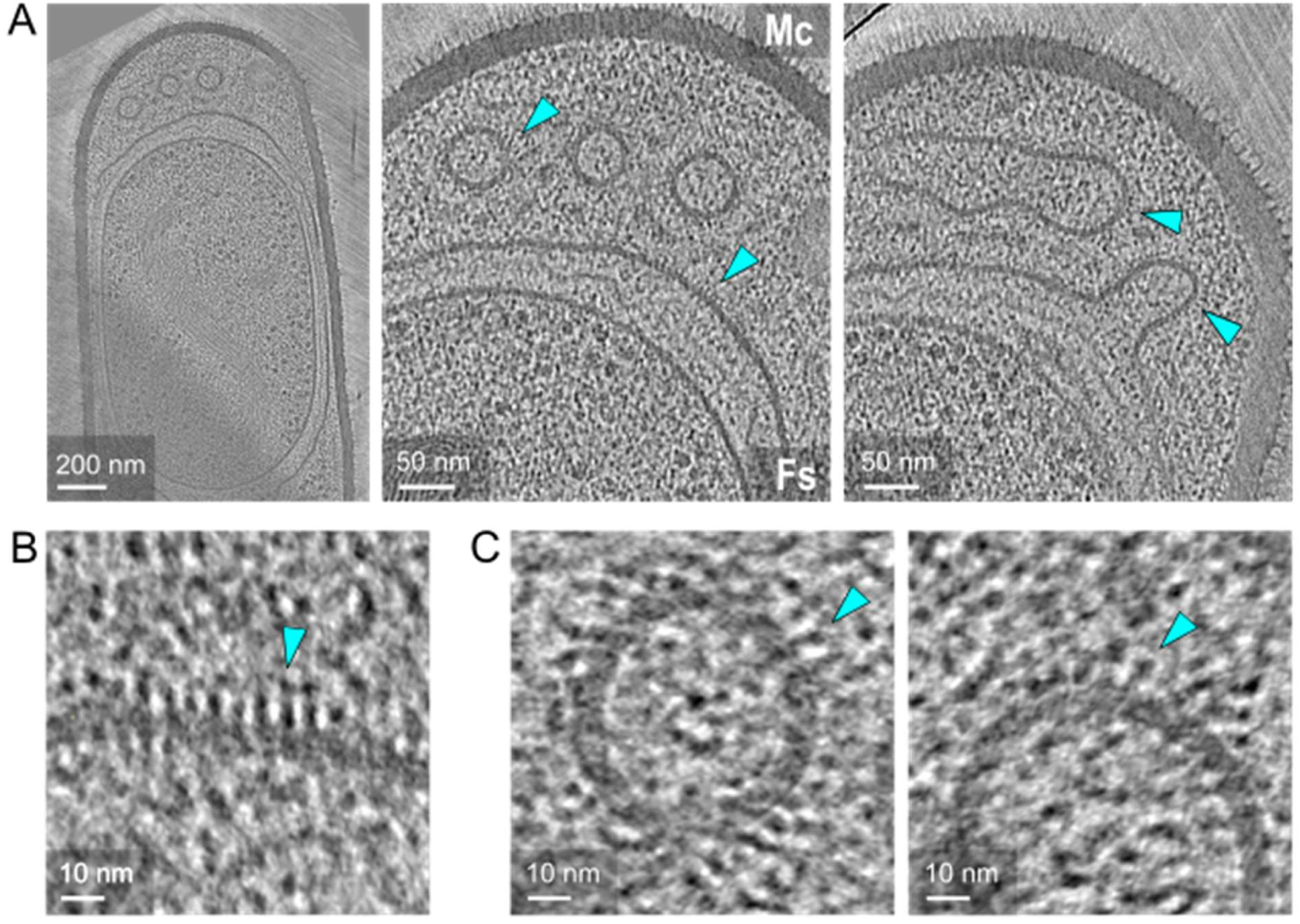
Localization of SpoIVA particles at the surface of positively curved membranes in *B. subtilis* sporangia. **A.** Slices through cryo-electron tomograms of *B. subtilis* sporangia harboring membrane defects, shown in full (**i**; scale bar, 200 nm) or magnified (**ii**; scale bars, 50 nm) views. The cyan inset highlights the zoomed area and the cyan arrowheads indicate SpoIVA particles. The tomogram was reconstructed from data acquired at medium magnification (pixel size 4.5 Å, high defocus). **B-C.** Zoomed-in regions from panel **A** highlighting SpoIVA particles at the surface of the outer forespore membrane (**B**), or along the convex face of membrane vesicles (**C**). Scale bars, 10 nm.

**Extended data Figure 2.**
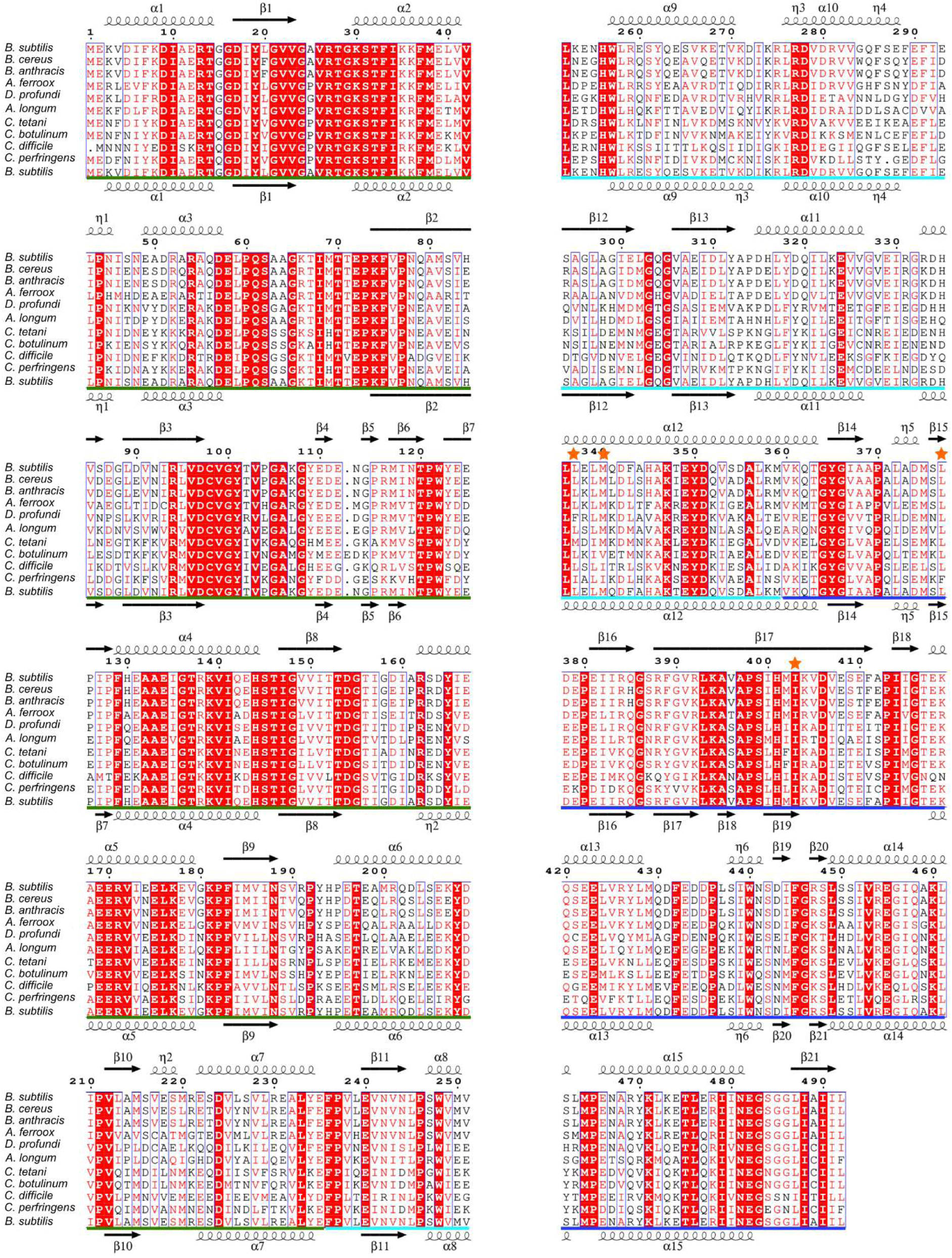
Conservation and secondary structure prediction of SpoIVA orthologues. A. Multiple sequence alignment of SpoIVA from *B. subtilis* with orthologues from *Bacillus cereus* (88% identity), *Bacillus anthracis* (88%), *Acidibacillus ferrooxidans* (*A. ferroox*; 71%), *Desulforamulus profundi* (60%), *Acetonema longum* (56%), *Clostridium tetani* (54%), *Clostridium botulinum* (53%), *Clostridium difficile* (48%), and *Clostridium perfringens* (48%). Conserved residues are highlighted in red boxes, while similar residues are shown as red letters in blue boxes. The predicted secondary structures of the SpoIVA dimer and monomer from *B. subtilis* are displayed above and below the alignment, respectively. The ATPase domain, the stalk and base regions are underlined in green, cyan and blue, respectively. The mutated interface residues are indicated by orange stars.

**Extended data Figure 3.**
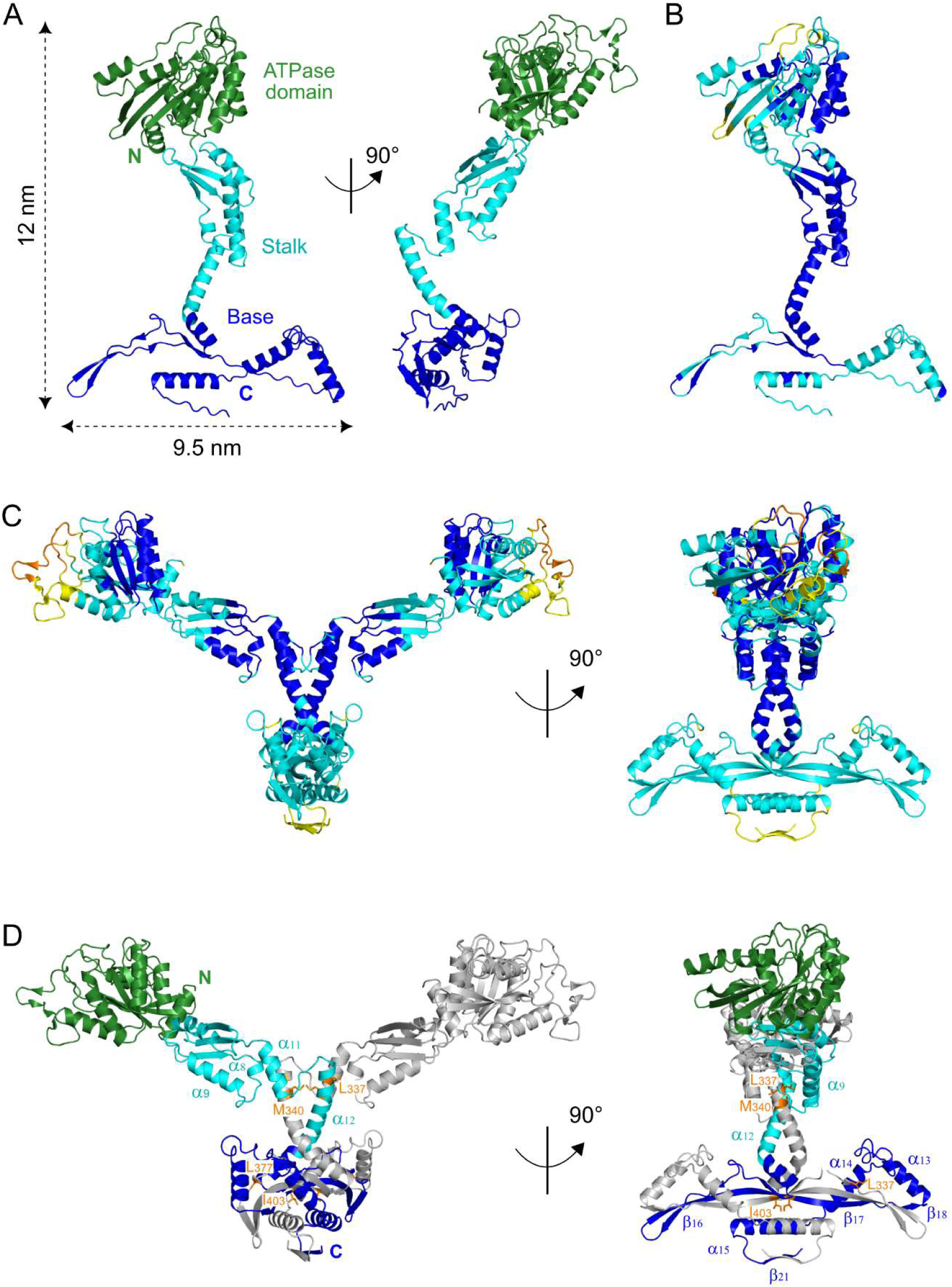
AlphaFold3 models of SpoIVA monomer and dimer. **A.** Two orthogonal views of the AlphaFold3-predicted SpoIVA monomer, highlighting the N-terminal globular ATPase domain (green), the stalk region (cyan), and the base region (blue). The protein dimensions are indicated. **B-C.** AlphaFold3-predicted SpoIVA monomer (**B**) and dimer (**C**) colored according to the pLDDT value. Dark blue, pLDDT > 90; light blue, 90 > pLDDT > 70; yellow, 70 > pLDDT > 50; orange, plDDT < 50. **D.** Two orthogonal views of the AlphaFold3-predicted SpoIVA dimer. One protomer is colored as in panel **A**, while the second protomer is shown in grey. Key secondary structures, mutated residues and positions of the N- and C-termini are annotated.

**Extended data Figure 4.**
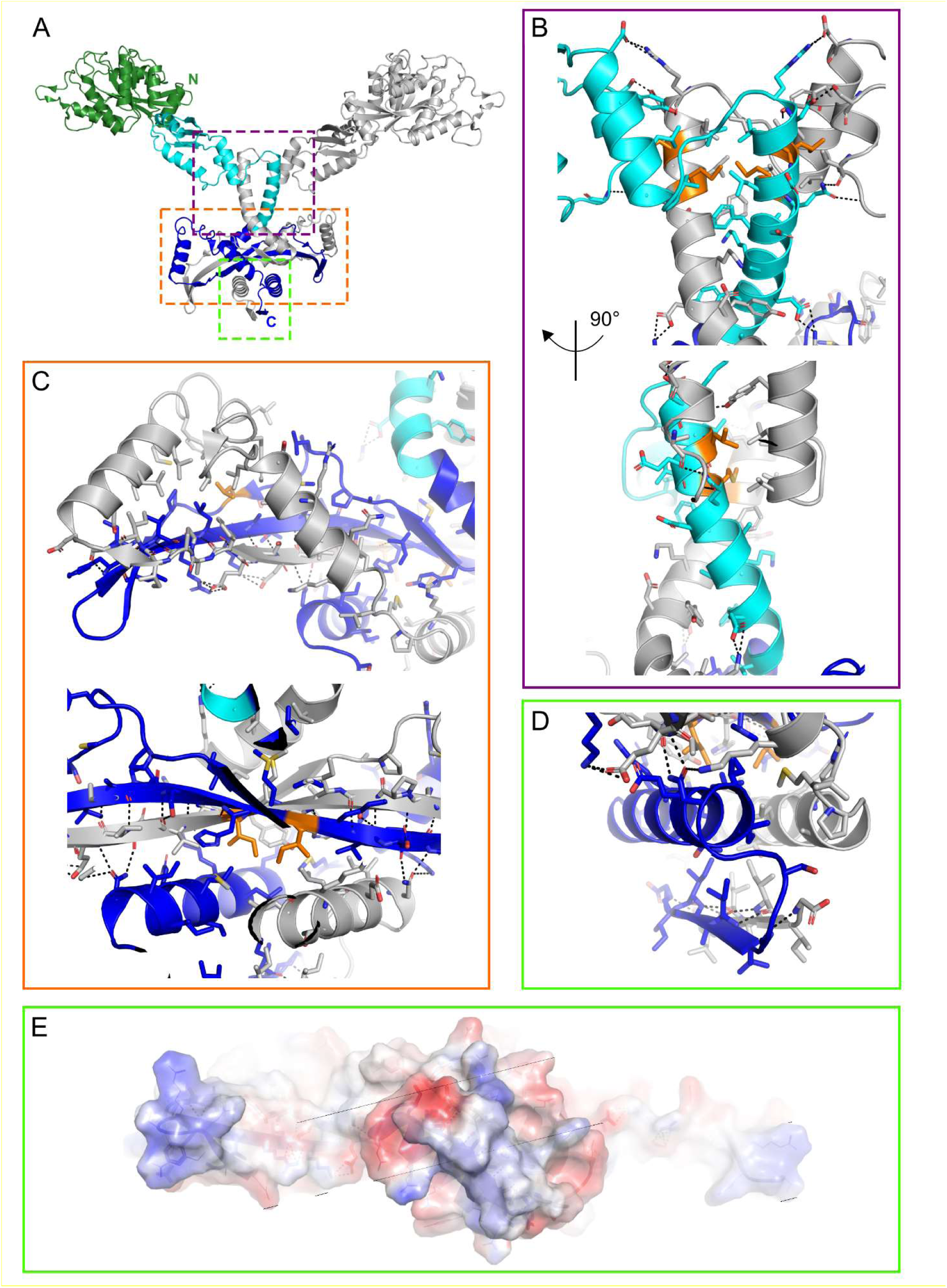
Structural prediction of the SpoIVA dimerization interface. **A**. Ribbon representation of the AlphaFold3-predicted SpoIVA homodimer. One protomer is colored in grey, while the ATPase domain (green), stalk (cyan) and base (blue) regions of the second protomer are highlighted. Dashed boxes indicate the regions magnified in panels **B-E**. **B-D.** Close-up views of the molecular interfaces established by the stalk (**B**), the upper base (**C**) featuring a long composite β-sheet, and the lower base (**D**), predominantly composed of hydrophobic residues. Interface residues are depicted in stick representation, with mutated residues colored in orange, and polar contacts shown as black dashed lines. **E.** Surface electrostatic potential of the lower base region, with negatively charged zones in red, positively charged zones in blue, and hydrophobic zones in white.

**Extended data Figure 5.**
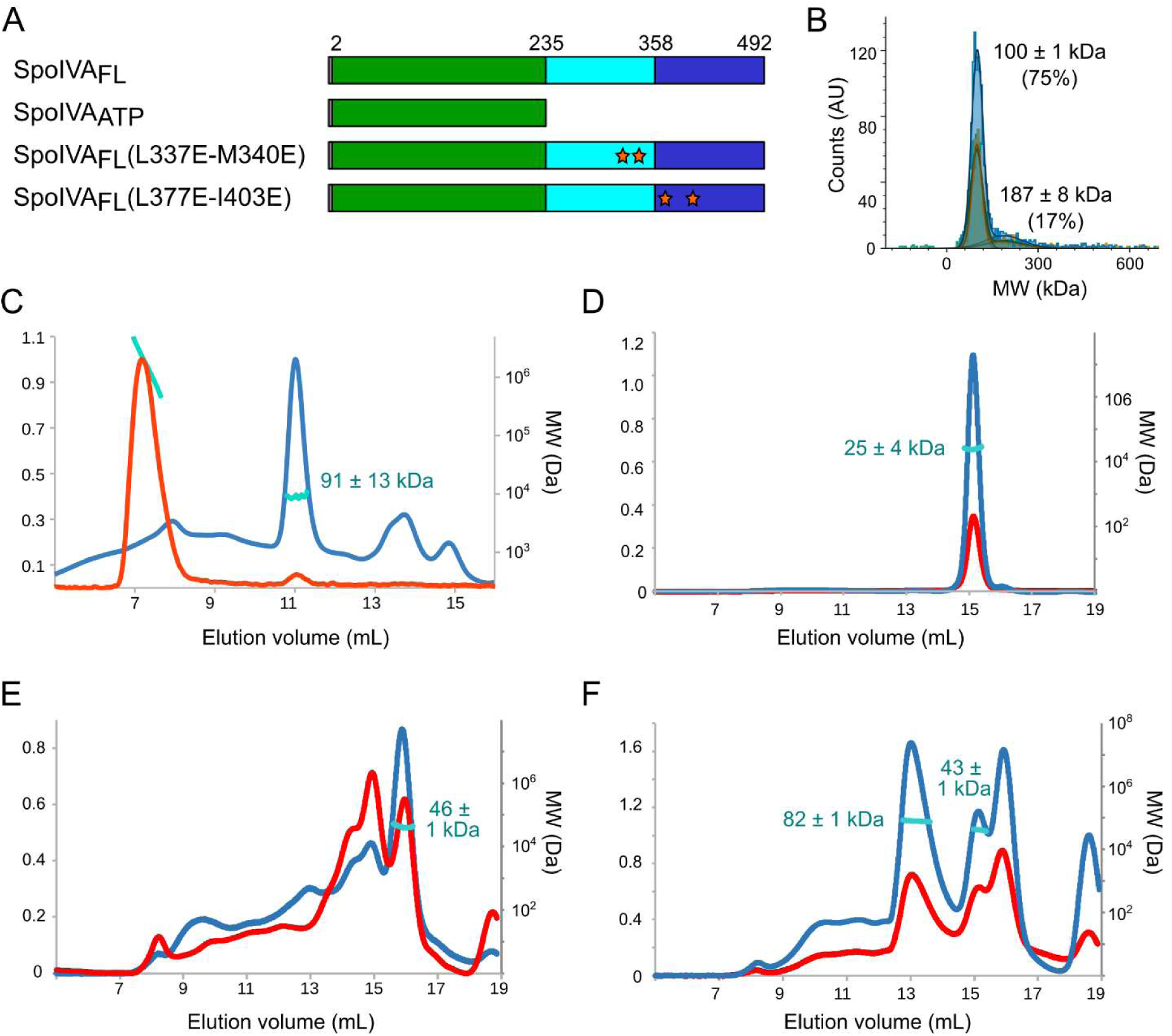
SpoIVA dimerization involves its stalk and base regions. **A.** Schematics showing the boundaries of the recombinant constructs and the position of the mutated interface residues (orange stars). **B.** Analysis of SpoIVA_FL_ by mass photometry. Counts in arbitrary units were plotted against the molecular weight (MW) in kDa, and the proportion of each species is indicated. Mass photometry measurements were performed in triplicates. **C-F.** Analysis of SpoIVA_FL_ (**C**), SpoIVA_ATP_ (**D**), SpoIVA_FL_(L337E-M340E) (**E**) and SpoIVA_FL_(L377E-I403E) (**F**) by SEC-MALLS. The blue curves show UV absorbance at 280 nm, the red curves correspond to light scattering, and the apparent MW of the main species is indicated. The left axis shows the relative scale in arbitrary unit for UV and light scattering signals; the right axis shows a log representation of the molecular weight (MW) in Da. SEC-MALLS analyses were performed in duplicates.

**Extended data Figure 6.**
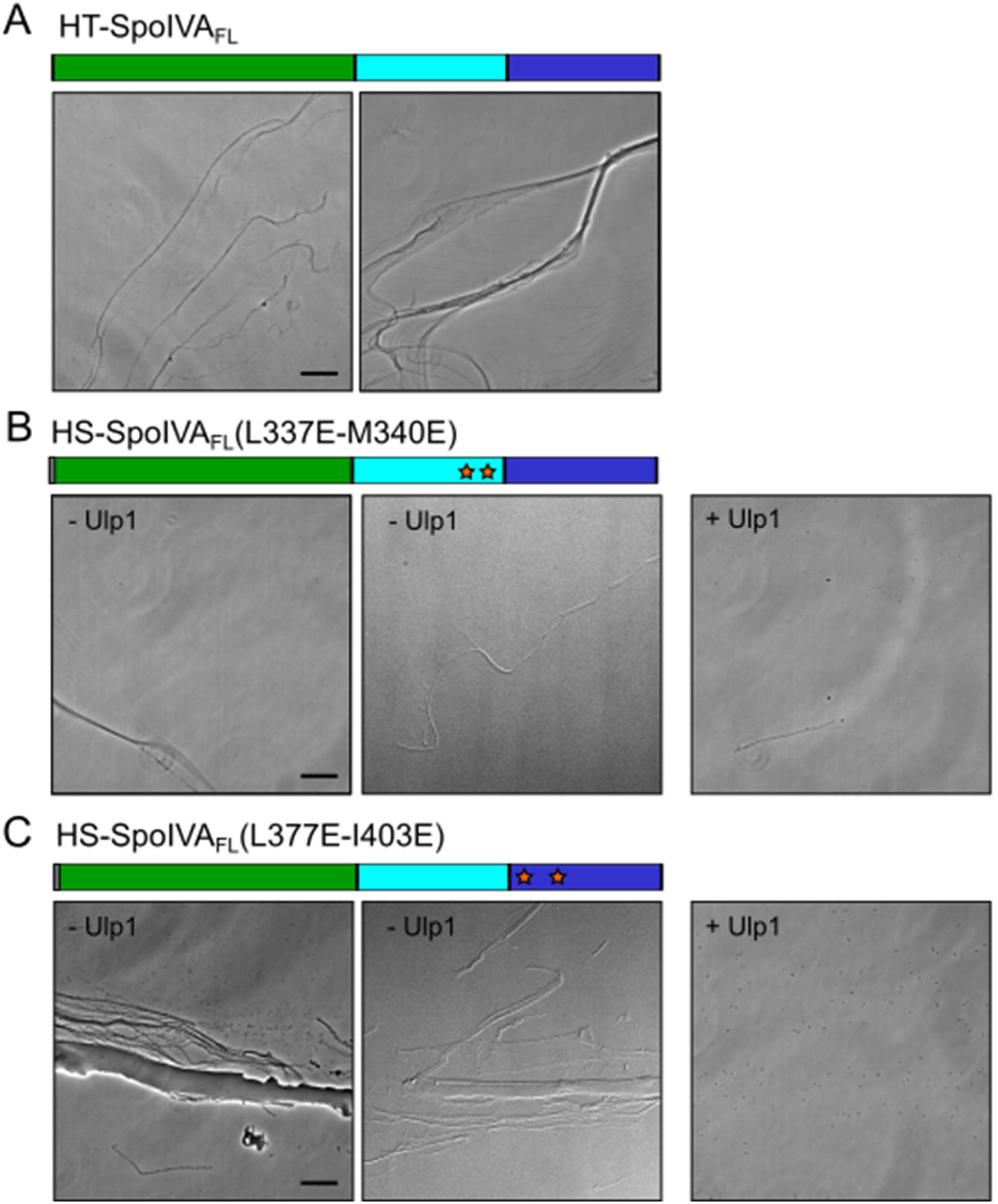
SpoIVA dimerization and filament formation. **A-C.** Phase-contrast images of purified His_6_-Thr-SpoIVA_FL_ (HT-SpoIVA_FL_, residues M1 to L492) (**A**), His_6_-SUMO-SpoIVA_FL_(L337E-M340E) (**B**) and His_6_-SUMO-SpoIVA _FL_(L377E-I403E) (**C**) variants after overnight incubation at 37 °C in the presence of 4 mM ATP, with or without HS-tag cleavage by the Ulp1 protease. Schematics illustrate the recombinant constructs used, with the mutated interface residues indicated by orange stars. Scale bar, 200 nm.

**Extended data Figure 7.**
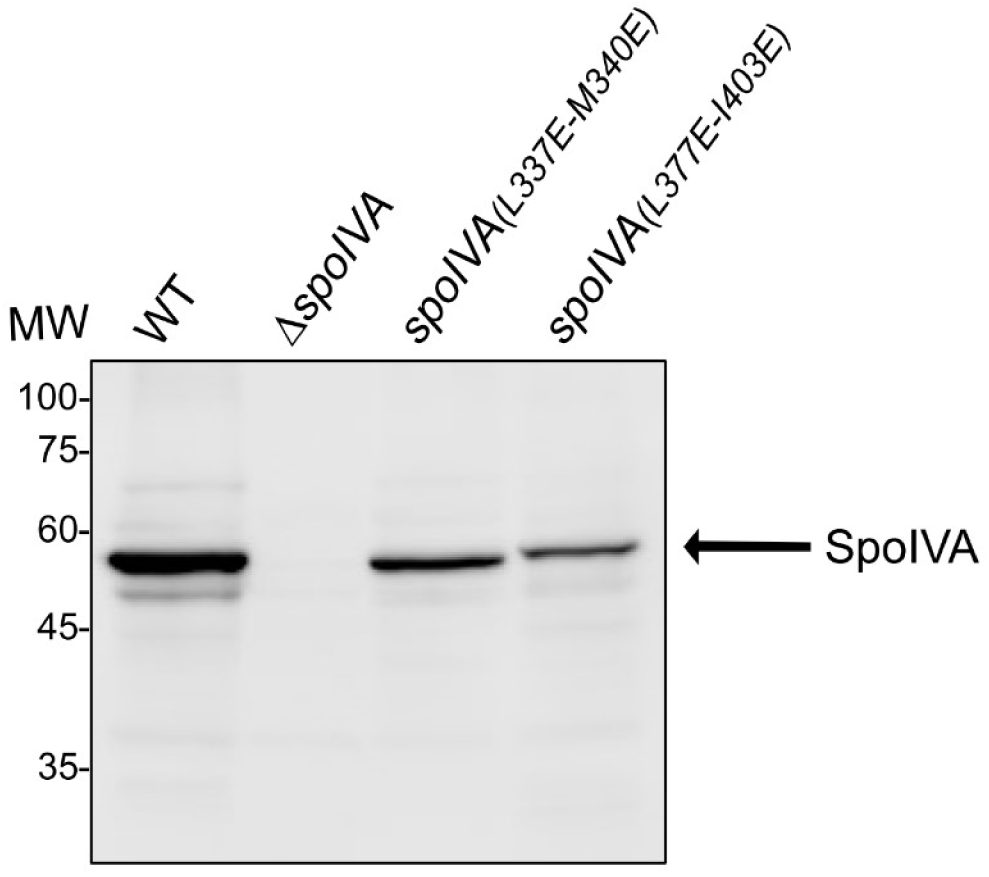
Western blot analysis of SpoIVA expression and stability. Western immunoblot of whole-cell lysates of wild-type (WT), *spoIVA(L337E-M340E)* and *spoIVA(L377E-I403E)* strains using anti-SpoIVA serum. The expected SpoIVA band is indicated by the black arrow. Molecular weight (MW) markers are indicated on the left.

**Extended data Figure 8.**
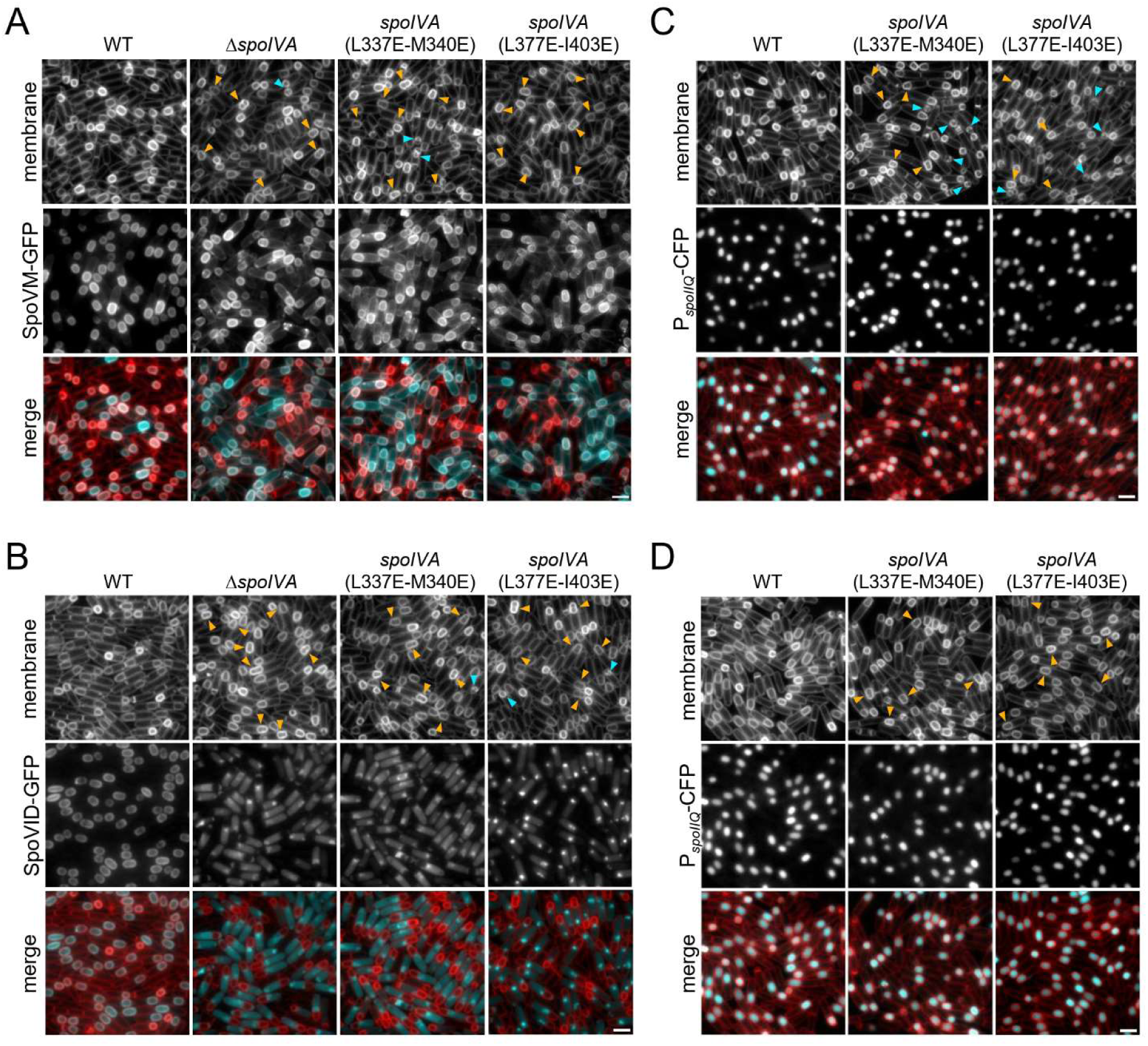
Disruption of SpoIVA dimerization alters its function. **A-B.** Fluorescence microscopy showing the localization of SpoVID-GFP (**C**) or SpoVM-GFP (**D**) in wild-type (WT), *spoIVA(L337E-M340E)*, or *spoIVA(L377E-I403E)* backgrounds at 3.5 h after the onset of sporulation. The GFP signals are false-coloured cyan in the merged images. Cell membranes were visualized with the TMA-DPH fluorescent dye, and are false-coloured red in the merged images. **C-D.** Sporulation progression in wild-type (WT), *spoIVA(L337E-M340E)*, or *spoIVA(L377E-I403E)* backgrounds at 2.5 h **(A)** and 3.5 h (**B**) after the onset of sporulation. The forespore cytoplasm was visualised using a PspoIIQ-cfp reporter, false-coloured cyan in merged images. Scale bars, 2 µm.

**Extended data Figure 9.**
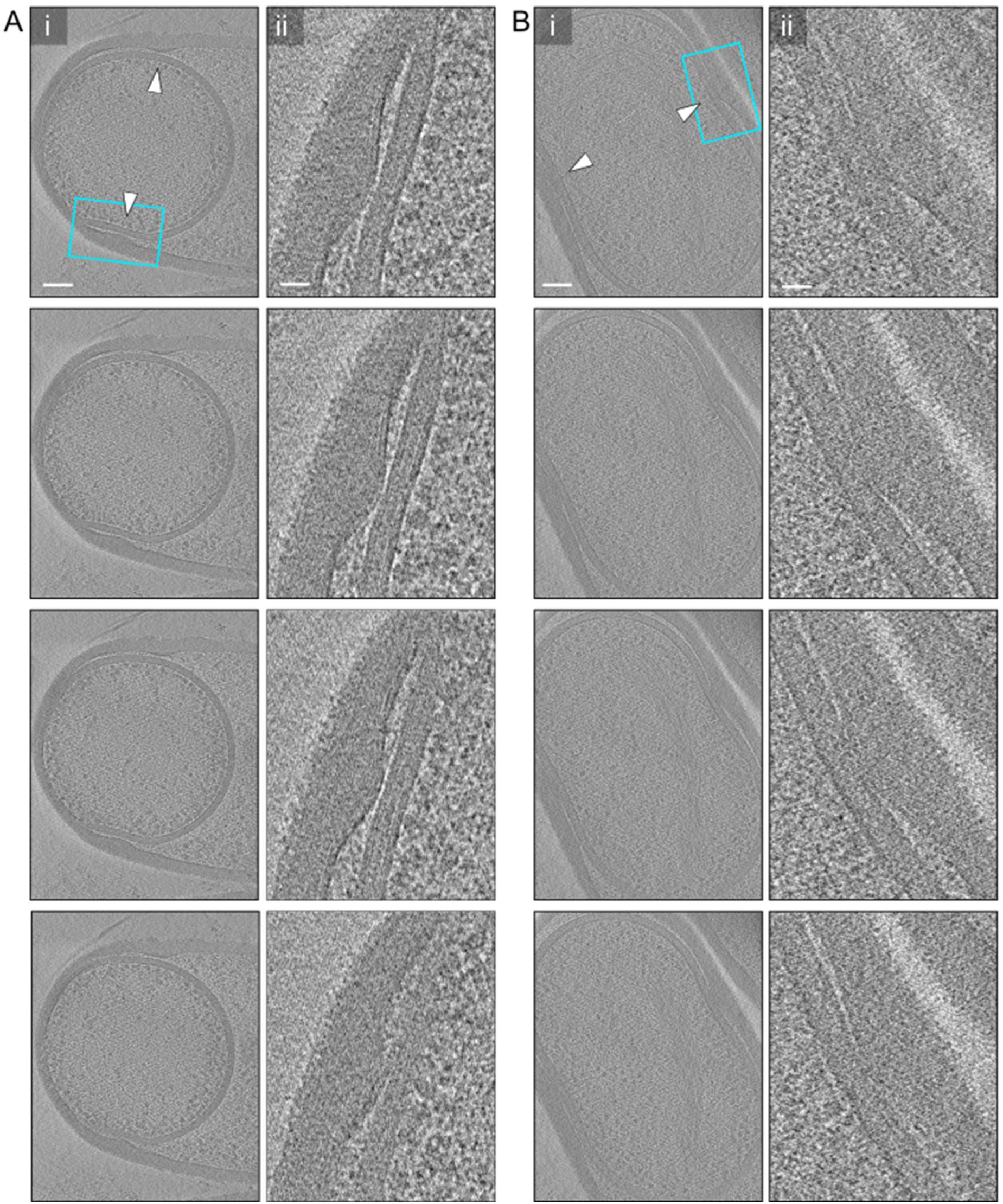
**A-B.** Slices through cryo-electron tomograms of engulfing (**A**) or engulfed (**B**) Δ*spoIVA* sporangia, presented in full views (**i**; scale bars, 100 nm) and magnified views (**ii**; scale bars, 25 nm) at incremental depths (top to bottom panels). The cyan inset highlights the zoomed area. Arrowheads indicate indentation events, where mother cell wall material compresses the forespore.

**Extended data Figure 10.**
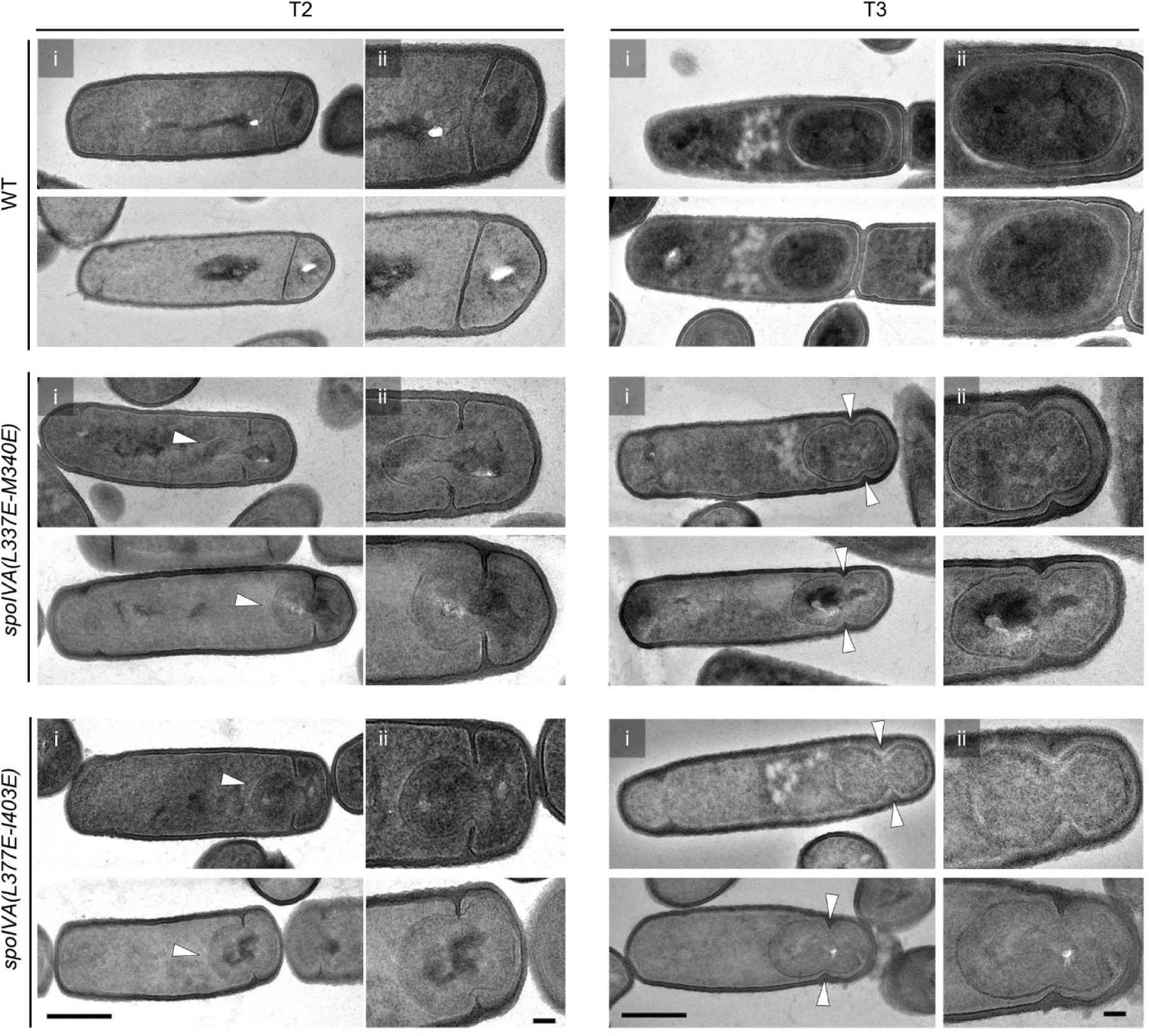
Full views (**i**, scale bar = 500 nm) and zooms (**ii**, scale bar = 100 nm) of transmission electron micrographs of resin-embedded sections of wild-type (WT), *spoIVA(L337E-M340E)* and *spoIVA(L377E-I403)* cells collected at 2 h (T2) and 3 h (T3) after the onset of sporulation. SpoIVA mutants bulging phenotype at T2 (single arrowheads) and indented phenotype (double arrowheads) after engulfment completion at T3.

**Extended Table 1.**
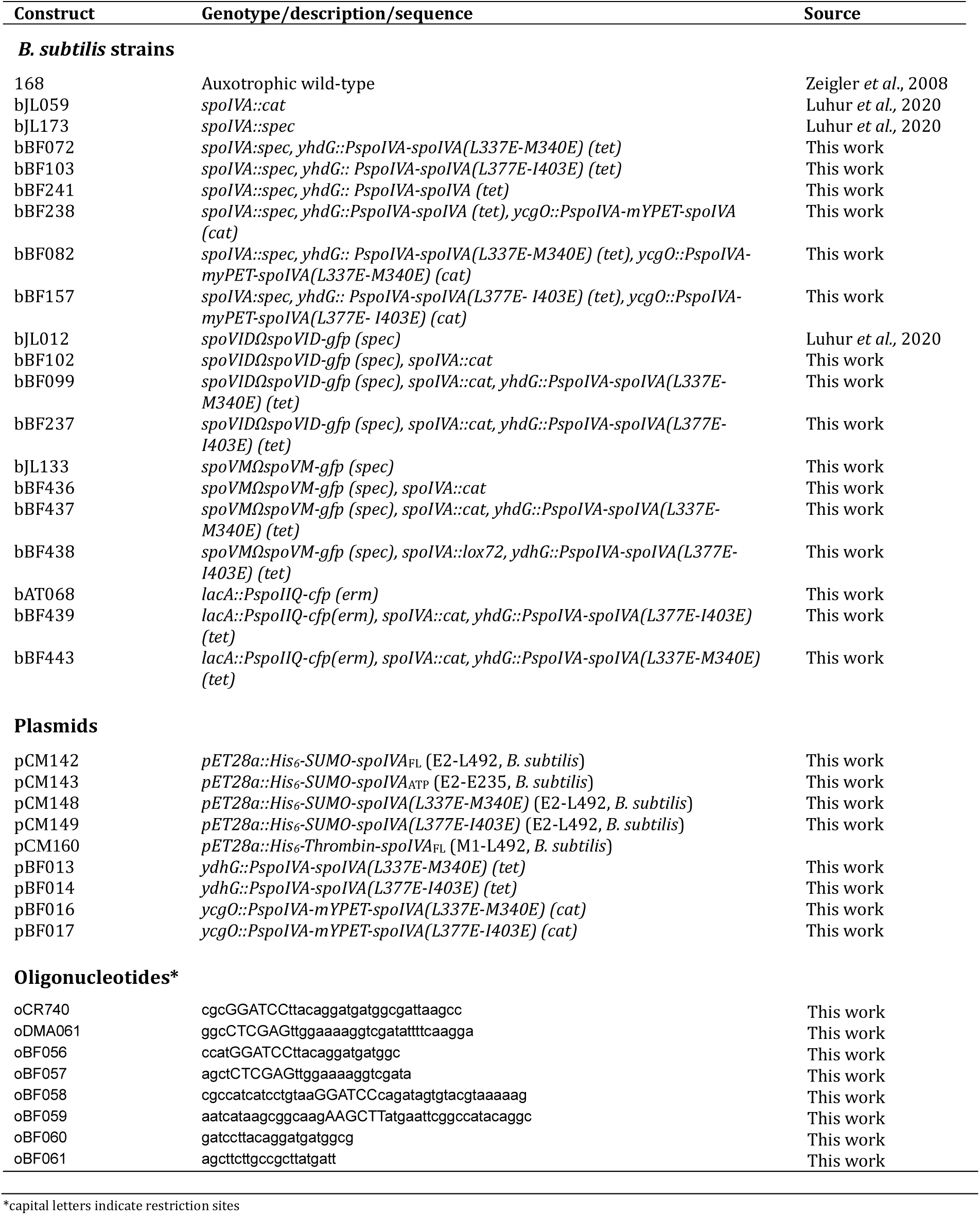
Strains, plasmids and oligonucleotides used in this study.

### Plasmid Construction

The HS-SpoIVA expression plasmids were synthesized by the GeneCust company (https://genecust.org). The gene fragments encoding SpoIVA from *B. subtilis* 168 were optimized for expression in *Escherichia coli*, and cloned between B*amH*I and *Xho*I in pET28a to be fused to a N-terminal His_6_-Thrombin-SUMO tag.

**pDMÅ06 [yhdG::P*spoIVA*-*spoIVA* (tet)]** was generated by a two-way ligation of a PCR product containing P*spoIVA*-*spoIVA* (amplified with oligonucleotide primers oCR740 and oDMÅ61) into pBB281 cut with *BamH*I*-Xho*I. pBB281 is an ectopic integration vector for recombination into *yhdG* locus (B. Burton and D.Z. Rudner, unpublished).

**pBF013 [*yhdG-PspoIVA-spoIVA-L337E-M340E (tet)*]** was generated by isothermal assembly joining a *Hind*III-*Bam*HI gene block (IDT) containing the relevant segment of *spoIVA* (*spoIVA* promoter and open-reading frame containing the specified mutations) and the pBB281 [*yhdG::tet*] backbone. The vector backbone was amplified to overlap with the gene block (using primers oBF058 and oBF059). pBB281 is an ectopic integration vector that facilitates recombination into the *yhdG* locus (B. Burton and D.Z. Rudner, unpublished).

**pBF014 [*yhdG::PspoIVA-spoIVA-L377E-I403E (tet*)]** was generated by isothermal assembly joining a *Hind*III-*Bam*HI gene block (IDT) containing the relevant segment of *spoIVA* (*spoIVA* promoter and open-reading frame containing the specified mutations) and pBB281 [*yhdG::tet*] backbone. The vector backbone was amplified to overlap with the gene block (using primers oBF058 and oBF059). pBB281 is an ectopic integration vector that facilitates recombination into the *yhdG* locus (B. Burton and D.Z. Rudner, unpublished).

**pBF016 *[ycgO::PspoIVA-mYPET-spoIVA-LL337E-M340E (cat)*]** was generated by a two-way ligation of a PCR product containing the *spoIVA-L377E-I403E* (amplified with oligonucleotide primers oBF056 and oBF057 and the gene block used to build pBF013 as template) into pJL006 cut with *Xho*I-*Bam*HI. pJL006 is an ectopic integration vector for recombination into the *ycgO* locus (Luhur et al., 2020).

**pBF017 [*ycgO::PspoIVA-mYPET-spoIVA-L377E-I403E (cat*)]** was generated by a two-way ligation of a PCR product containing the *spoIVA-L377E-I403E* (amplified with oligonucleotide primers oBF056 and oBF057, and the gene block used to build pBF014 as template) into pJL006 cut with *Xho*I-*Bam*HI. pJL006 is an ectopic integration vector for recombination into the *ycgO* locus (Luhur et al., 2020).

